# cAMP controls a trafficking mechanism that directs the neuron specificity and subcellular placement of electrical synapses

**DOI:** 10.1101/2021.05.12.443836

**Authors:** Sierra Palumbos, Rachel Skelton, Rebecca McWhirter, Amanda Mitchell, Isaiah Swann, Sydney Heifner, Steve Von Stetina, David M Miller

## Abstract

Electrical synapses are established between specific neurons and within distinct subcellular compartments, but the mechanisms that direct gap junction assembly in the nervous system are largely unknown. Here we show that a transcriptional program tunes cAMP signaling to direct the neuron-specific assembly and placement of electrical synapses in the *C. elegans* motor circuit. For these studies, we use live cell imaging to visualize electrical synapses in vivo and a novel optogenetic assay to confirm that they are functional. In VA motor neurons, the UNC-4 transcription factor blocks expression of cAMP antagonists that promote gap junction miswiring. In *unc-4* mutants, VA electrical synapses are established with an alternative synaptic partner and are repositioned from the VA axon to soma. We show that cAMP counters these effects by driving gap junction trafficking into the VA axon for electrical synapse assembly. Thus, our experiments in an intact nervous system establish that cAMP regulates gap junction trafficking for the biogenesis of electrical synapses.

## INTRODUCTION

Neuronal function depends on the neuron-specific assembly of both chemical and electrical synapses. In comparison to chemical synapses (Margeta and Shen, 2010; Margeta et al., 2008; Sanes and Yamagata, 2009; Williams et al., 2010), strikingly little is known of the pathways that direct the formation of electrical synapses between specific neurons (Hendi et al., 2019; Hestrin and Galarreta, 2005). This disparity is significant as electrical synapses account for nearly 20% of connections in mature nervous systems (Cook et al., 2019; Kim et al., 2012; White et al., 1986) and are required for diverse neural processes (Allen et al., 2011; Kawano et al., 2011; Miller et al., 1992; Otopalik et al., 2019; Phelan et al., 1998; Song et al., 2016; Walker and Schafer, 2020; Zolnik and Connors, 2016). In addition, the differential placement of electrical synapses in specific subcellular compartments (e.g. axon vs. soma) strongly influences neuronal output (Tamás et al., 2000; Wang et al., 2017). Thus, a deeper understanding of mechanisms that direct assembly of electrical synapses in specific circuits and to the correct subcellular location is of fundamental importance to developmental neuroscience.

Electrical synapses, or gap junctions, mediate the rapid transfer of ions and small molecules, thus coupling the membrane potentials of connected neurons (Hormuzdi et al., 2004; Meda and Spray, 2000; Pereda et al., 2013). In mammals and other vertebrates, gap junctions are assembled from connexins which oligomerize to form hexagonal hemichannels in each neuron (Maeda et al., 2009; Sosinsky and Nicholson, 2005). Hemichannels in adjacent neurons appose one another to form a gap junction array. In an intriguing example of convergent evolution, the invertebrate gap junction subunits, innexins, adopt strikingly similar topology and function as connexins despite lacking clear sequence homology (Skerrett and Williams, 2016). The likelihood of conserved mechanisms for gap junction assembly, however, is supported by the findings that innexin-containing gap junctions can assemble and function in vertebrate cells (Dykes et al., 2004; Phelan et al., 1998; Starich et al., 2009), and conversely, that connexins can form functional gap junctions when expressed in invertebrate cells (Meng and Yan, 2020; Rabinowitch et al., 2014). It follows that invertebrate model organisms, with their powerful genetic tools and readily-accessible live-cell imaging, can be exploited to identify regulators of gap junction biogenesis that may also specify electrical synapses in mammalian neurons (Meng and Yan, 2020; Meng et al., 2016; Schneider et al., 2012; Von Stetina et al., 2007).

Early experiments in cultured cells suggested a straightforward model in which neuron-specific electrical synapses arise *solely* from differential expression of gap junction subunits. This idea derived from the observation that overexpression of either connexins or innexins is sufficient to drive assembly of gap junctions at the interface between random pairs of adjacent cells (Elfgang et al., 1995; Rabinowitch et al., 2014; Teubner et al., 2000). Investigations of this question *in vivo*, however, have revealed that assembly of neuron-specific electrical synapses requires additional regulatory mechanisms. For example, in many instances, gap junctions do not assemble between adjacent neurons that express compatible gap junction subunits (Bhattacharya et al., 2019; Fukuda, 2017; Greb et al., 2017; Miller et al., 1992; Von Stetina et al., 2005; Yao et al., 2016). Thus, developmental programs that direct trafficking and assembly of gap junction components are likely necessary for the placement of electrical synapses between specific neurons.

A wide range of molecular regulators of gap junction homeostasis have been defined by experiments in cultured cells. For example, specific protein kinases modulate gap junction function and trafficking (Cooper and Lampe, 2002; Dunn et al., 2012; Lampe and Lau, 2004; Pidoux and Taskén, 2015; Solan and Lampe, 2005, 2009, 2016). Notably, Protein Kinase A (PKA), a 3’,5’-cyclic adenosine monophosphate (cAMP) dependent kinase, can promote gap junction coupling, potentially by regulating of trafficking of Cx43 connexons to the plasma membrane (Atkinson et al., 1995; Burghardt et al., 1995; Holm et al., 1999; Lampe et al., 2001; Matsumura et al., 2006; Ouyang et al., 2005; Paulson et al., 2000; TenBroek et al., 2001; Thevenin et al., 2013). Few studies, however, have asked whether similar mechanisms regulate gap junction placement in an intact nervous system.

Here, we exploit the simplicity and accessibility of the *C. elegans* nervous system to identify components that direct the formation of neuron-specific gap junctions. We used a genetic screen to identify two antagonists of cAMP signaling, a phosphodiesterase (PDE-1) and a GPCR (FRPR-17), that regulate the neuron-specificity and subcellular placement of gap junctions in the motor circuit. Both *pde-1* and *frpr-17* are normally turned off in Ventral A Class (VA) motor neurons by the UNC-4 transcription factor. Ectopic expression of PDE-1 and FRPR-17 in an *unc-4* mutant, however, results in the creation of electrical synapses between VAs and a new interneuron partner and also repositions gap junctions from the VA axon to the cell soma. Thus, UNC-4 regulates cAMP signaling to control both the neuron specificity and subcellular placement of electrical synapses in the motor circuit. We performed genetic and pharmacologic experiments to confirm that cAMP regulates assembly of functional electrical synapses. Our findings show that elevated cAMP promotes trafficking of gap junction components from the VA cell soma for assembly of electrical synapses in the VA axon. Our results suggest that this trafficking mechanism facilitates the creation of additional neuron-specific electrical synapses as VA neurons expand in size during larval development. In contrast, when cAMP levels are reduced, gap junction components fail to exit the VA cell soma where they accumulate and form ectopic electrical synapses with a different interneuron partner. These results demonstrate that cAMP regulates a trafficking mechanism that controls both neuron specificity and subcellular placement of electrical synapses in an intact nervous system.

## RESULTS

### UNC-4-regulated genes direct gap junction specificity

In the *C. elegans* motor circuit, a transcriptional program directs synaptic specificity of the VA motor neurons. VA motor neurons adopt both chemical and electrical synapses with the interneuron AVA (VA→AVA), whereas VBs, the sister cells of VAs, exclusively establish electrical synapses with the interneuron AVB (VB→AVB) (Cook et al., 2019; White et al., 1986). These connections are required to drive either backward (VA→AVA) or forward (VB→AVB) locomotion (Chalfie et al., 1985; Kawano et al., 2011; Wen et al., 2012). The transcription factor UNC-4 is selectively expressed in VA motor neurons where it controls synaptic specificity; *unc-4* mutants are unable to move backward because chemical and electrical synapses with AVA are replaced with ectopic electrical input from AVB (VA→AVB) (**Figure 1A-C**). In addition to the misassembly of VA electrical synapses with AVB, VA gap junctions are also repositioned from the VA axon to the cell soma in *unc-4* mutants (**Figure 1A**) (Miller and Niemeyer, 1995; Schneider et al., 2012; Von Stetina et al., 2007; White et al., 1992). Thus, UNC-4 regulates both the neuron specificity and subcellular placement of electrical synapses in VA neurons. To identify the molecular players that influence these effects, we used a combination of cell-specific RNA-Seq profiling and genetic tests to detect transcripts regulated by UNC-4 that control the neuron-specific assembly of electrical synapses.

**Figure 1:**
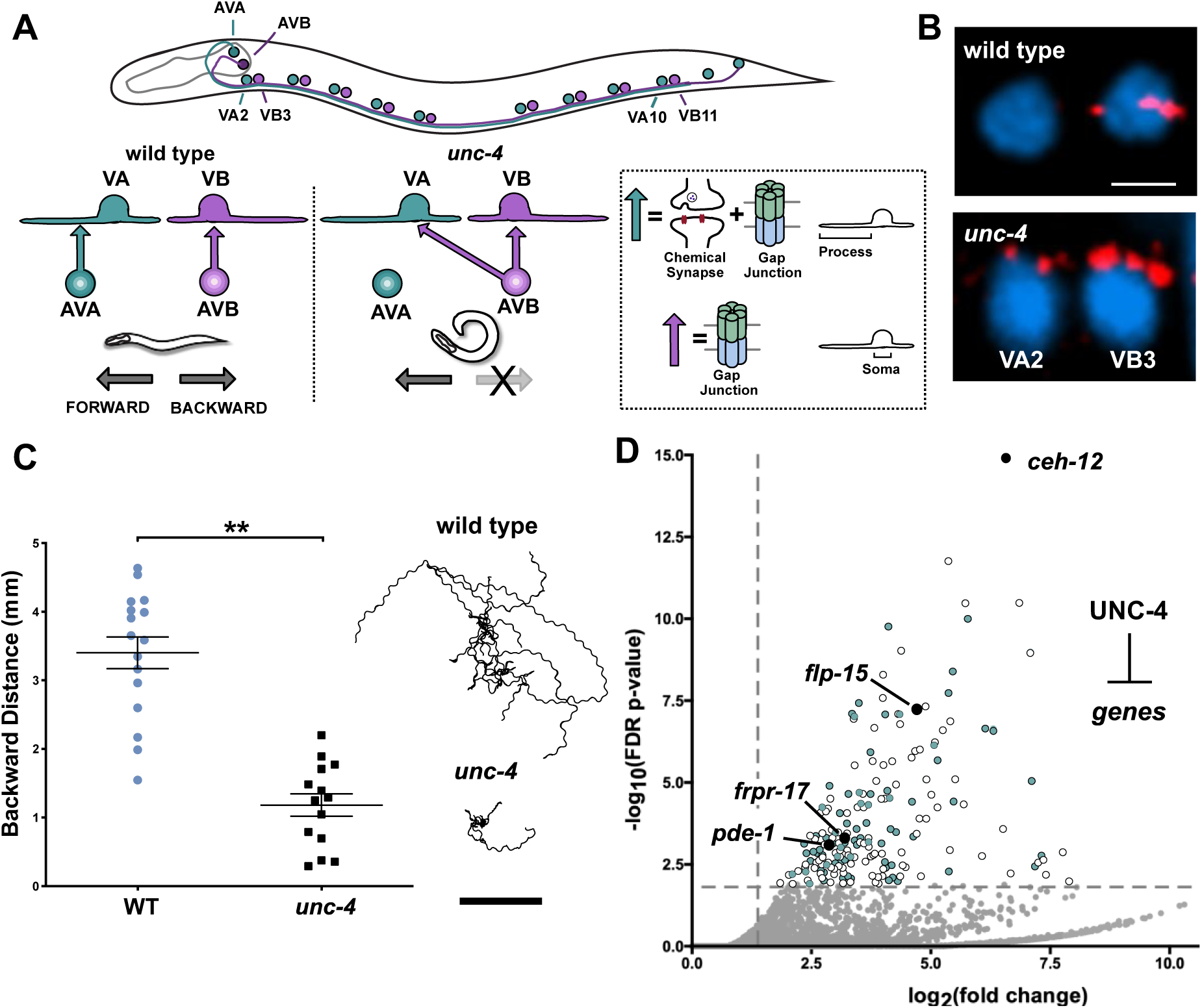
UNC-4 functions as a transcriptional repressor to direct gap junction specificity. **A.** Schematic of motor circuit regulated by UNC-4. AVA and AVB interneurons in the head region extend axons into the ventral nerve cord to synapse with VA and VB motor neurons. In the wild type, AVA establishes chemical and electrical synapses with VA motor neuron process (VA→AVA). AVB assembles electrical synapses with VB cell soma (VB→AVB). *unc-4* mutants are miswired with AVB electrical synapses on VA cell soma (VA→AVB) **B.** Representative images of gap junction wiring in wild type (top) and the hypomorphic allele *unc-4(e2323)* (bottom). AVB gap junctions are labeled with UNC-7s::GFP (red) and VA2 and VB3 soma are stained with DAPI (blue). In *unc-4* mutants, UNC-7s::GFP puncta are readily detected on the VA2 cell soma and correspond to ectopic VA→AVB (bottom). Scale bar = 2.5µm. **C.** (Left) Quantification of backward distance traveled by wild type (light blue) (n = 15) and *unc-4(e2323)* (black) (n = 13) in a 3-minute period. Student’s t-test, ** = p< .01. Data are mean +/- SE. (Right) Representative tracks of 10 wild type (top) and *unc- 4(e2323)* (bottom) L4 worms in a 3-minute period. Scale bar = 2mm. **D.** Volcano plot of upregulated transcripts (> 2X, FDR p-value < .01) detected in *unc-4(e120)* null mutant VAs (see Methods). 80/214 upregulated targets were tested by RNAi or in genetic mutants for suppression of the Unc-4 movement defect (blue dots). Four genes (black) were identified as suppressors of a hypomorphic *unc-4* mutant (*e2323* or *e2322*).

We used an intersectional labeling strategy for FACS-isolation of VA neurons from L2 larval stage animals, the developmental period in which *unc-4* function is required (Miller et al., 1992; Spencer et al., 2014). RNA-seq analysis identified >2000 transcripts that are significantly elevated in wild-type VAs in comparison to all L2 larval stage cells. Robust expression of known VA cell marker genes (e.g., *unc-4, unc-3, del-1, cfi-1)* in this data set validates FACS enrichment of VA motor neurons (**Figure S1)** (Kerk et al., 2017; Miller and Niemeyer, 1995; Von Stetina et al., 2007; Taylor et al., 2020). Next, we compared the RNA-Seq profiles of wild-type vs. *unc-4* mutant VAs. We detected over 300 transcripts (98 down-regulated, 214 up-regulated) as differentially expressed in *unc-4* mutant VAs (**Figure S2**). We focused on the 214 transcripts that are up-regulated in *unc-4* mutant VAs because UNC-4 normally functions as a transcriptional repressor (Miller et al., 1992; Pflugrad et al., 1997; Von Stetina et al., 2007; Winnier et al., 1999) (**Figure 1D**). The homeodomain transcription factor gene, *ceh-12,* was the most significantly upregulated transcript in *unc-4* mutant VAs. This finding is consistent with previous microarray and GFP-reporter results showing that UNC-4 blocks *ceh-12* expression (Schneider et al., 2012; Von Stetina et al., 2007). Importantly, ectopic expression of *ceh-12* in *unc-4* mutants is restricted to a subset of VA neurons in the posterior ventral nerve cord (VNC). Genetic epistasis experiments confirmed that ectopic *ceh-12* is regionally required for the gap junction miswiring defect of posterior *unc-4* mutant VAs (Von Stetina, 2007). The local effect of *ceh-12* on gap junction specificity predicts that UNC-4 must regulate other targets that act in anterior VAs to sustain the wild-type pattern of electrical connectivity.

To identify additional regulators of VA gap junction specificity, we next used RNAi and available loss-of-function mutants in a “suppressor” screen of target genes that are ectopically expressed in *unc-4* mutant VAs. Knockdown of an UNC-4 target that drives the Unc-4 miswiring defect is predicted to at least partially restore backward movement in an *unc-4* mutant, an outcome we describe as Unc-4 suppression (**Figure S2**). For this assay, we limited our analysis to the 214 transcripts that are upregulated in *unc-4* mutant VAs. We tested 80 genes with corresponding mammalian homologs and detected three “suppressor” loci (*pde-1, frpr-17, flp-15)* that significantly improved locomotion in a sensitized *unc-4* mutant background (**Figure 2B-C, Figure S2, Table S1**). Notably, *pde-1* and *frpr-17* are both predicted to antagonize cAMP signaling (**Figure 2A**). PDE-1 is functionally homologous to the mammalian phosphodiesterase PDE1A (77.4%, BLAST e-value 3e-173) with calmodulin and calcium-dependent enzymatic activity that degrades both cAMP and cGMP (Lugnier, 2006). FRPR-17 corresponds to an orphan GPCR that is predicted to bind FMRF-type neuropeptides and couple to Gαi/o (99% coupling score, PredCouple2) (**Figure S2**). *C. elegans* expresses a single Gαi/o, GOA-1/Gαo, which is homologous to the mammalian GNAI2 (100%, BLAST e-value 0) and GNAO1 (82.2%, BLAST e-value 0) that antagonize (AC) Adenylyl Cyclase-dependent synthesis of cAMP (Muntean et al., 2021). Therefore, ectopic expression of FRPR-17 in *unc-4* mutant VAs could activate GOA-1 with a consequent reduction in cAMP. Taken together, these results suggest that UNC-4 may act to preserve cAMP signaling in VA neurons by blocking expression of both PDE-1, which degrades cAMP, and FRPR-17, which limits cAMP synthesis. Our findings also argue that cAMP signaling maintains VA inputs that are required for backward locomotion.

**Figure 2:**
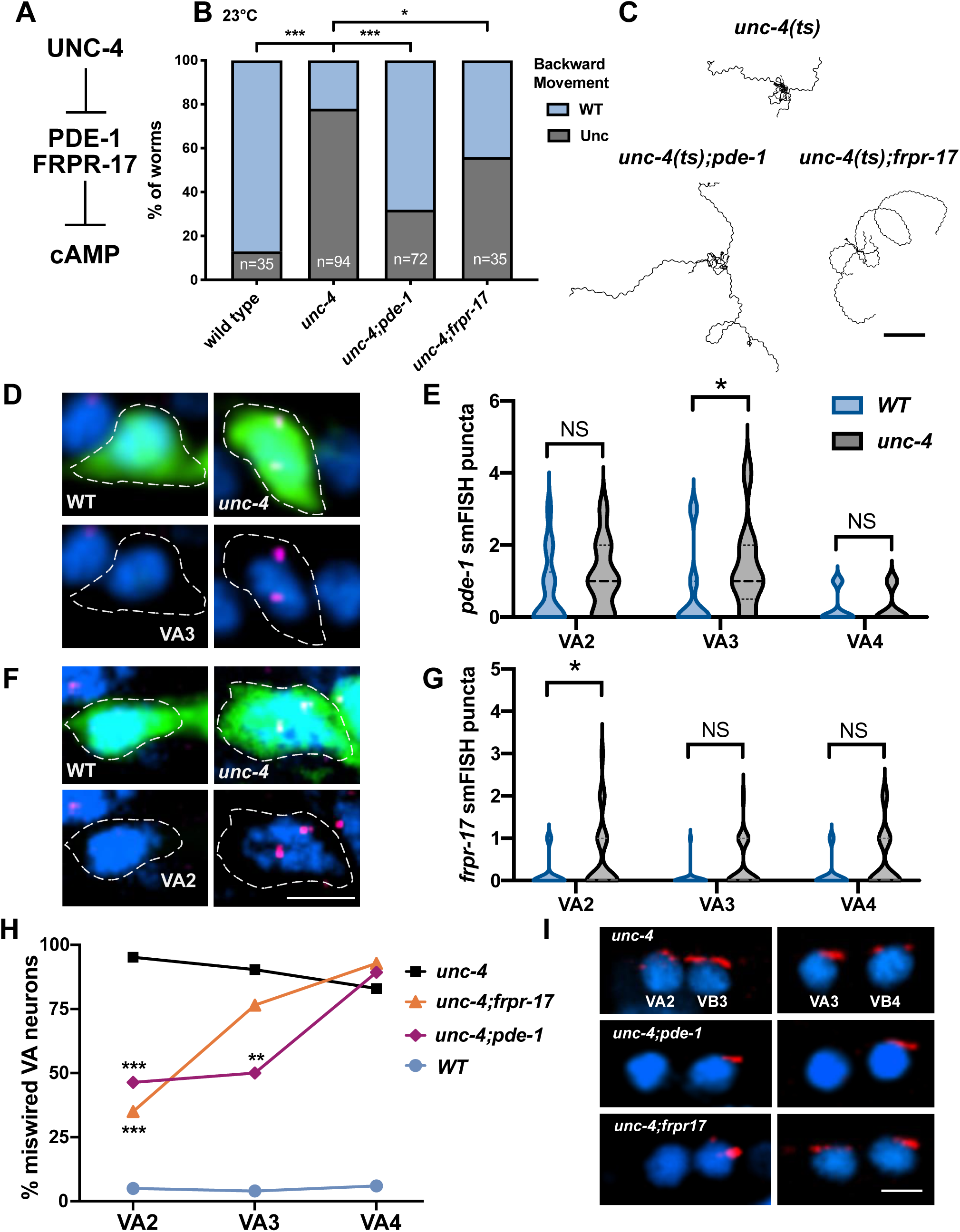
UNC-4 maintains cAMP through distinct mechanisms in different VA motor neurons. **A.** Model of UNC-4-dependent regulation of cAMP. UNC-4 blocks expression of PDE-1/Phosphodiesterase and FRPR-17/GPCR, both predicted antagonists of cAMP. **B.** Quantification of head tap resulting in backward movement scored as either wild type (WT) or uncoordinated (Unc). At 23°C, backward movement is strongly impaired in *unc-4(ts)* worms. Genetic ablation of either *pde-1* or *frpr-17* partially restores backward locomotion to *unc-4(ts).* Fisher’s Exact test versus *unc-4 (ts)*. * = p< 0.05, *** = p<0.001. **C.** Representative tracks of *unc-4(ts), unc-4(ts);pde-1* and *unc-4(ts);frpr-17* L4 worms at 23°C in a 3-minute period. Scale bar = 2mm. **D.** Representative images of VA3 in wild type (left) and *unc-4(e120)* (right). Dashed lines denote cell soma marked with GFP (top), labeled with DAPI (blue), and *pde-1* smFISH probe (magenta puncta) in L2 stage larvae. **E.** Quantification of *pde-1* smFISH puncta in VA motor neurons. Violin plots for *pde-1* smFISH puncta in WT (light blue) and *unc-4(e120)* (grey) in L2 stage VA neurons. Dashed line denotes median. Mann-Whitney test, * p = 0.037. N.S., Not Significant. n>20 per VA. **F.** Representative images of VA2 in wild type (left) and *unc-4(e120)* (right). Dashed lines denote cell soma marked with GFP (top) and labeled with DAPI (blue) and *frpr-17* smFISH probe (magenta puncta). **G.** Quantification of *frpr-17*smFISH puncta in VA motor neurons. Violin plots for *frpr-17* smFISH puncta in WT (light blue) and *unc-4*(*e120*) (grey) in L2 stage VA neurons. Dashed line denotes median. Mann-Whitney test, * p= 0.039. N.S., Not Significant. n>25 per VA. **H.** Quantification of ectopic VA→AVB gap junctions plotted as percent of each anterior VA neuron (VA2, VA3, VA4) in *unc-4(e2323)* (black), *unc-4(e2323);frpr-17* (orange)*, unc-4(e2323);pde-1* (magenta) and wild type (WT) (light blue). Fisher’s exact versus *unc-4*. N>15 for each VA. ** = p<0.01, *** = p<0.001. **I.** Representative images of UNC-7S::GFP marking AVB gap junctions (red) with VA and VB motor neurons in *unc-4(e2323)* (top) *unc-4(e2323);pde-1* (middle), and *unc-4(e2323);frpr-17* (top). DAPI (blue) labels VA and VB nuclei. Scale bar = 2.5 µm.

### UNC-4 regulates cAMP signaling through neuron-specific gene regulation

Our RNA-sequencing analysis indicates that *pde-1* (4.5X, FDR p<.01) and *frpr-17* (8.2X, FDR p<.01) are upregulated in *unc-4* mutant VAs (**Figure 1D**). As an independent test of this finding, we used single molecule fluorescent *in situ* hybridization (smFISH) to visualize *pde-1* and *frpr-17* transcripts in VA neurons. We focused on VAs in the anterior VNC (VA2, VA3, VA4) because we have previously shown that the UNC-4-regulated target, CEH-12, is ectopically expressed in posterior VA neurons (VA8, VA9, VA10) to drive miswiring in *unc-4* mutants. In wild-type worms, we detected *pde-1* smFISH puncta in VA2 and VA3. In *unc-4* mutants, we observed a significant increase in *pde-1* transcripts in VA3 (**Figure 2D-E**). *pde-1* is also ectopically expressed in a subset of posterior VA neurons in *unc-4* mutants (data not shown). In contrast, *frpr-17* was selectively elevated in *unc-4* mutant VA2 with little *frpr-17* transcript detected in wild-type VA2-4 (**Figure 2F-G**). These findings are consistent with our RNA-seq results showing that UNC-4 negatively regulates *frpr-17* and *pde-1* expression in VA neurons. Interestingly, UNC-4 appears to modulate cAMP levels by regulating different targets in specific VAs, e.g. *frpr-17* in VA2 and *pde-1* in VA3.

### UNC-4 blocks the formation of ectopic electrical synapses through differential neuron-specific gene regulation

As noted above, genetic knockdown of either *pde-1* or *frpr-17* partially suppressed the Unc-4 movement defect (**Figure 2B, C**). These findings argue that ectopic expression of *pde-1* and *frpr-17* contributes to the impaired backward movement of *unc-4* mutants. Because the Unc-4 movement phenotype is also correlated with the formation of ectopic VA→AVB gap junctions (**Figure 1A, B**), we next asked if *pde-1* and *frpr-17* are necessary for the VA→AVB wiring defect in *unc-4* mutants. For this assay, we expressed the GFP- tagged innexin UNC-7 (UNC-7S::GFP) in the interneuron AVB to detect ectopic gap junctions with VA neurons (VA→AVB) (Starich et al., 2009; Von Stetina et al., 2007). The specificity of these electrical synapses is readily detectable because AVB gap junctions are characteristically positioned on motor neuron cell soma (White et al., 1986, 1992). Thus, AVB gap junctions can be determined by scoring the co-localization of UNC-7S::GFP puncta with the nuclei of ventral cord motor neurons marked with the DNA- specific dye, DAPI (4′,6-diamidino-2-phenylindole) (**Figure 1B**) (Schneider et al., 2012; Von Stetina et al., 2007). If ectopic *pde-1* expression is required for the Unc-4 miswiring defect (i.e., VA→AVB electrical synapses), then the occurrence of VA→AVB gap junctions should be reduced in *unc-4; pde-1* double mutants compared to *unc-4* alone. We observed significantly fewer VA→AVB gap junctions in *unc-4; pde-1* mutants for VA2 and VA3 but not for VA4 (**Figure 2 H-I**). This finding confirms that elevated PDE-1/phosphodiesterase activity in *unc-4* mutants results in the formation of ectopic VA→AVB electrical synapses in VA2 and VA3. Our smFISH assay showed that *pde-1* transcripts are detectable in VA2 and elevated in VA3 in *unc-4* mutants, suggesting that *pde-1* expression in both VA2 and VA3 favors miswiring.

We also used the UNC-7S::GFP marker to show that VA→AVB gap junctions are reduced in VA2 in *unc-4; frpr-17* double mutants, but not in VA3 or in VA4 (**Figure 2H-I**). This selective effect of the *frpr-17* mutation on VA2 wiring is consistent with our smFISH results showing that the *frpr-17* transcript is exclusively detected in *unc-4* mutant VA2 neurons in the anterior nerve cord (**Figure 2F-G**). We note that our detection of electrical synapses for individual neurons (e.g., VA2 vs VA3), highlights the utility of the *C. elegans* nervous system for the investigation of mechanisms that regulate the neuron-specificity of gap junction assembly. Together, our findings point to a model in which UNC-4 antagonizes the formation of ectopic VA→AVB gap junctions by blocking expression of different target genes in specific VA neurons (e.g., *frpr-17* in VA2 and *pde-1* in VA3) that antagonize cAMP signaling.

### UNC-4 preserves cAMP to direct the neuron specificity of electrical synapses

Having shown that UNC-4 downregulates expression of two presumptive antagonists of cAMP signaling (PDE-1 and FRPR-17) to regulate the neuron specificity of gap junction assembly, we hypothesized that elevated cAMP signaling prevents the formation of ectopic VA→AVB electrical synapses. In that case, other pharmacological or genetic manipulations that elevate cAMP should similarly restore backward locomotion to an *unc-4* mutant. As an initial test of this prediction, we utilized the cell-permeable, non-hydrolyzable cAMP analog, 8-Bromo-cAMP (8-Br-cAMP) to phenocopy global elevation of cAMP (Hussey et al., 2017). *unc-4* mutant worms fed 8- Bromo-cAMP exhibited significant restoration of backward locomotion (**Figure 3B**). This result is consistent with the hypothesis that cAMP levels are depleted in *unc-4* mutants and that reduced cAMP signaling is sufficient to disrupt wiring of the backward movement circuit.

**Figure 3:**
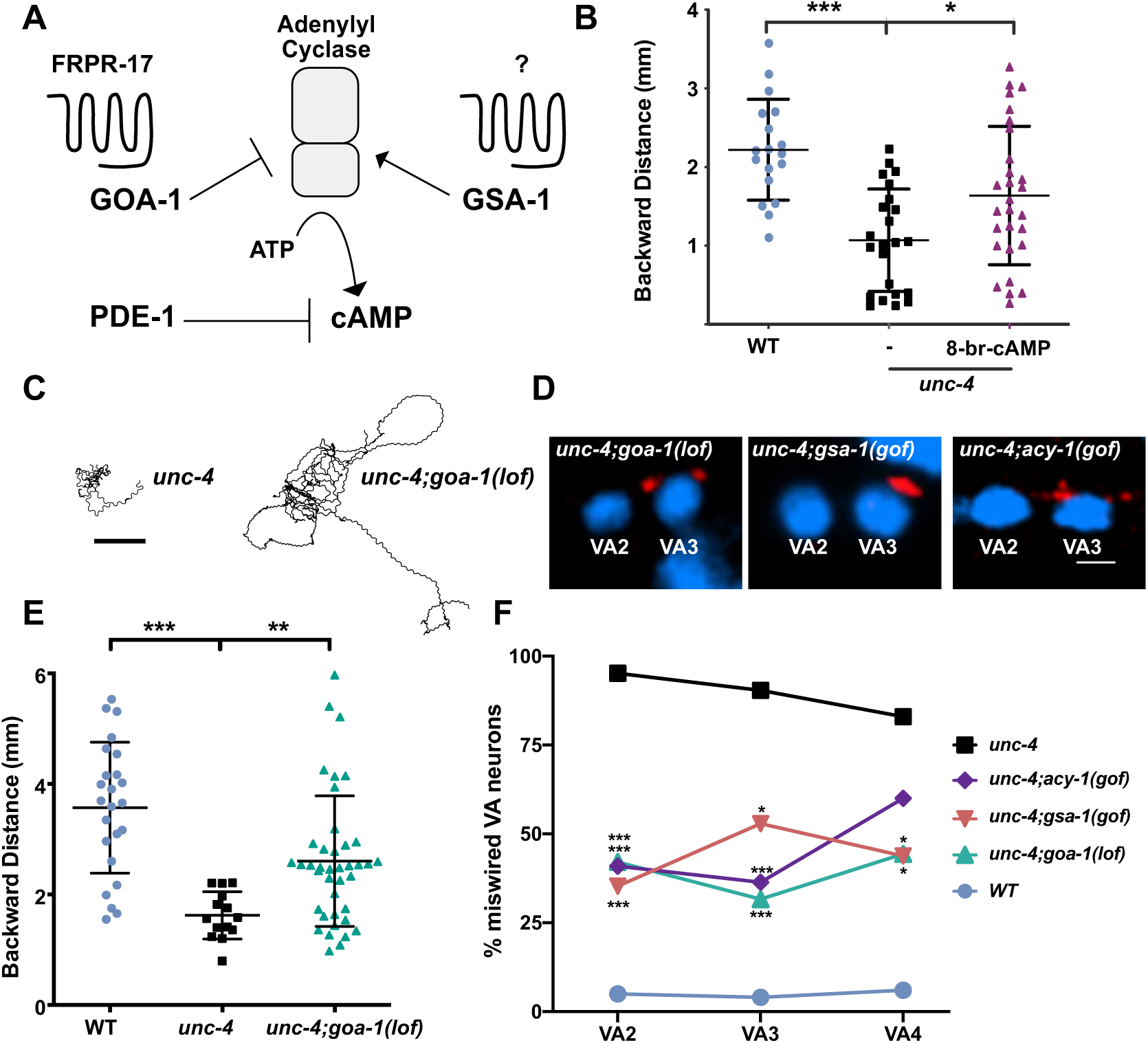
cAMP maintains electrical synapses for backward locomotion. **A.** Schematic representation of cAMP regulation. The G-protein GSA-1/GaS promotes Adenylyl Cyclase-dependent synthesis of cAMP whereas GOA-1/GaO antagonizes cAMP production. The GPCR, FRPR-17, is predicted to couple to GOA-1. PDE-1/phosphodiesterase degrades cAMP. **B.** Quantification of backward distance traveled in a 3-minute period of wild type (light blue), *unc-4* (black), and *unc-4* + 8-Br-cAMP (maroon). Growth on 8-Br-cAMP partially restores backward locomotion to *unc-4(e2323)* mutant animals. One-way ANOVA, N > 15 for each genotype. * p=.0169, **** = p<.0001. **C.** Representative tracks of ten *unc-4(2323)* (left) and ten *unc-4(e2323);goa-1(lof)* (right) L4 larvae in a 3-minute period. Scale bar = 2mm. **D.** Representative images of UNC-7S::GFP marking AVB gap junctions with VA and VB motor neurons in *unc-4(e2323);goa-1(lof), unc-4(e2323);gsa-1(gof),* and *unc-4(e2323);acy-1(gof).* DAPI (blue) labels VA and VB nuclei. Scale bar = 5 µm. **E.** Quantification of backward distance traveled by wild type (light blue), *unc-4(e2323)* (black), and *unc-4(e2323);goa-1(lof)* (green). *unc-4(e2323);goa-1(lof)* double mutants show partial restoration of backward locomotion vs *unc-4*. One-way ANOVA, N > 15 for each genotype, ** p=.0089, **** = p<.0001. **F.** Quantification of ectopic VA→AVB gap junctions marked with UNC-7S::GFP. Data are shown as percent of miswired VA motor neurons (VA2, VA3, VA4) in wild type, *unc-4*, and in double mutants *unc-4(e2323);goa-1(lof), unc-4(e2323);gsa-1(gof),* and *unc-4(e2323);acy-1(gof*). Fisher’s exact test vs. *unc-4*, n > 15 for each VA, * = p<.05, ***= p<.001. lof = loss-of-function, gof = gain-of-function.

To further delineate the role of cAMP in electrical synaptic specificity, we used a series of genetic approaches to elevate cAMP levels *in vivo*. In *C. elegans*, the biosynthetic enzyme adenylyl cyclase (ACY-1/AC) is regulated by antagonistic G-protein pathways that either stimulate (GSA-1/GαS) or reduce (GOA-1/GαO) cAMP production (**Figure 3A**) (Govindan et al., 2006). Therefore, activation of either ACY-1/AC or GSA-1/GαS should increase cAMP. cAMP levels should also be elevated by genetic ablation of GOA-1/GαO, which normally impedes production of cAMP by adenylate cyclase. First, we asked if the loss of GOA-1/GαO function could suppress the Unc-4 movement defect. We determined that backward movement in *unc-4; goa-1(lof)* mutants was significantly improved in comparison to *unc-4* (**Figure 3C-D, Video S1**). This result is consistent with the hypothesis that elevated cAMP promotes VA connections that drive backward movement. As a direct test of this idea, we used the UNC-7S::GFP marker to score ectopic VA→AVB gap junctions in *unc-4; goa-1(lof)* double mutants and confirmed a significant reduction of VA→AVB miswiring in comparison to *unc-4* mutant VAs (**Figure 3E-F**). Strikingly, the VA→AVB miswiring defect was suppressed in all three anterior VAs (VA2, VA3, VA4) in *unc-4; goa-1 (lof)* animals. We have also observed that VA→AVB miswiring is similarly suppressed in multiple posteriorly located VAs in *unc-4; goa-1(lof)* (data not shown). Thus, these observations suggest that the specificity of VA gap junction formation is globally sensitive to cAMP signaling.

Next, we utilized a hyperactive allele of GSA-1/GαS to elevate cAMP. In this case, we again observed that ectopic VA→AVB gap junctions are reduced in VA2-VA4 (**Figure 3F**). As both GSA-1 and GOA-1 also affect cholinergic signaling, we considered the possibility that excess ACh release from VA motor neurons could be sufficient to account for suppression of the VA→AVB gap junction miswiring defect in *unc-4*; *goa-1(lof)* and *unc-4; gsa-1(gof)* mutants. However, genetic experiments with downstream components involved in cholinergic signaling did not alter VA gap junction specificity **(Figure S3)**. Thus, our findings favor a model in which GOA-1/Gαo and GSA-1/Gαs regulation of cAMP signaling, and not ACh release, mediates the specificity of electrical synapses in this circuit. This idea is also supported by our finding that a gain-of-function adenylate cyclase allele, *acy-1(gof),* which increases production of cAMP, suppresses the Unc-4 gap junction miswiring defect in VA2-VA3 (**Figure 3D**). Taken together, these findings suggest that elevated cAMP antagonizes the formation of ectopic VA→AVB electrical synapses and that UNC-4 turns off specific target genes (*e.g., pde-1, frpr-17*) to preserve cAMP signaling in VA neurons.

### cAMP acts within a temporal developmental window to promote gap junction specificity

We next sought to identify the temporal window in which cAMP regulates neuron-specific gap junctions in VA neurons. Previous studies established that UNC-4 function is required in VAs during a defined period of larval development (L2-L3) or 10-20 hours post hatching (HPH) (Miller et al., 1992). We therefore predicted that elevated cAMP should also be necessary during this developmental window to establish VA inputs that drive backward movement. To test this idea, we used an optogenetic tool for temporal and tissue-specific elevation of cAMP. The Beggiatoa-photoactivated adenylyl cyclase (bPAC) produces cAMP in response to blue light (Steuer Costa et al., 2017) (**Figure 4A**). We used the *unc-17* promoter to drive bPAC (*Punc-17*::bPAC) expression in cholinergic neurons, including VAs, in *unc-4* mutant animals. We exposed different groups of *unc-4;pUnc-17::bPAC* animals to blue light for separate 10-hour periods during three developmental stages: (A) 0-10 HPH (B) 10-20 HPH (C) 20-30 HPH. For each treatment, backward locomotion was examined later, at the adult stage, to rule out potential acute effects of cAMP elevation on motor circuit function (Steuer Costa et al., 2017). These experiments revealed that photostimulation of bPAC during 10-20 HPH (Period B) resulted in improved backward locomotion in adult *unc-4;pUnc-17::bPAC* worms, whereas transient exposure to blue light at either an earlier (0-10 HPH) or later (10-20 HPH) developmental period did not enhance backward locomotion (**Figure 4B**). These findings are consistent with a model in which UNC-4 acts within a critical developmental window to preserve cAMP signaling and thus maintain VA function in the backward motor circuit.

**Figure 4:**
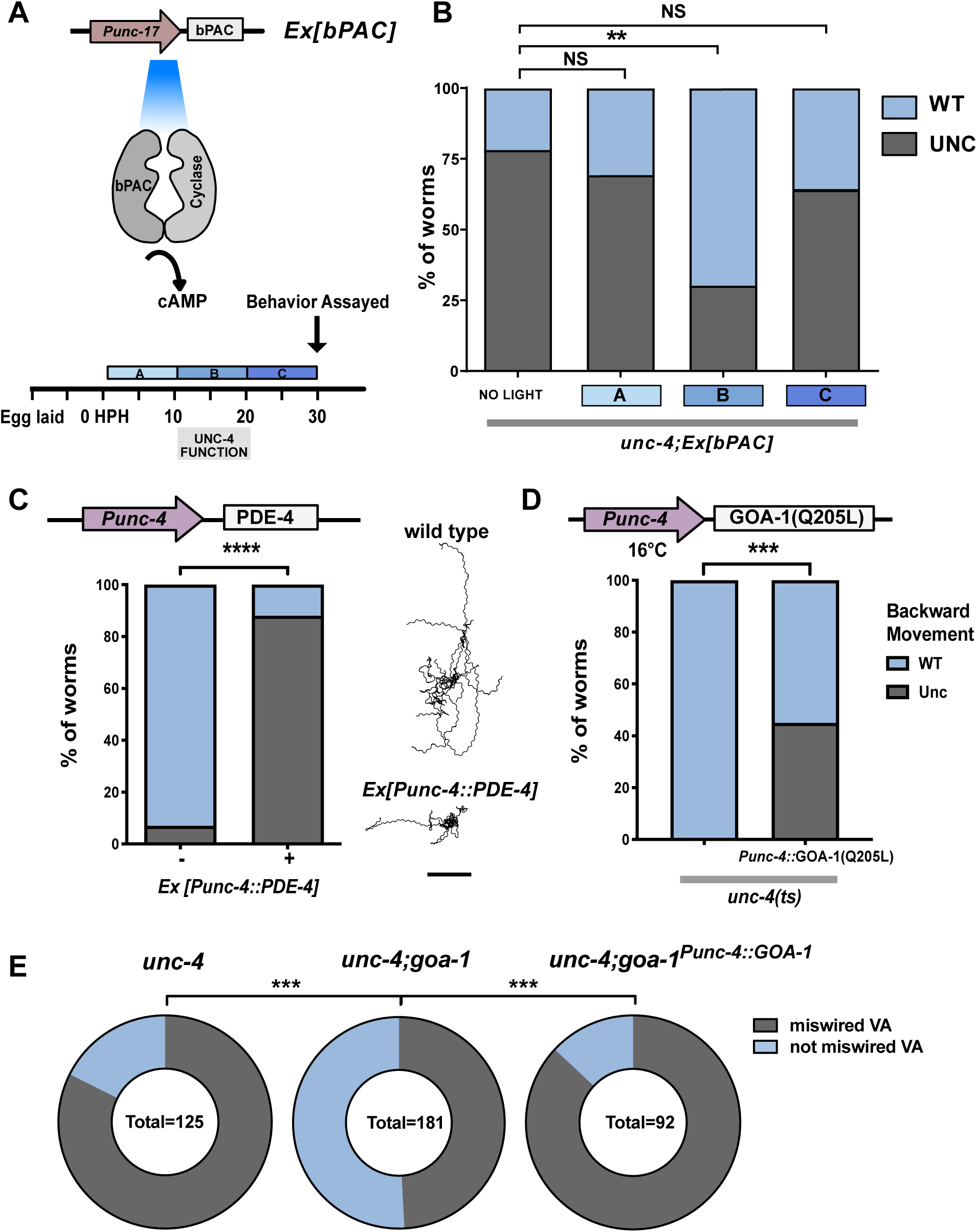
cAMP is required in VA neurons to maintain electrical synapses for backward locomotion. **A.** Schematic of strategy for optogenetic activation of cAMP synthesis. The promoter, *Punc-17*, drives expression of photoactivatable Adenylyl Cyclase (*bPAC*) in cholinergic neurons in *unc-4(e2323)* mutants. *unc-4(e2323); Ex[bPAC]* worms were exposed to blue light during three developmental windows (A) 0-10 HPH, (B) 10-20 HPH, and (C) 20-30 HPH. UNC-4 is required in developmental period (B) for wild-type backward movement (Miller et al., 1992). HPH = hours post hatching.**B.** Quantification of backward locomotion. All worms are *unc-4(e2323);Ex[Punc-17::bPAC]* and were exposed to blue light during one developmental window, (A), (B) or (C). Blue light activation during developmental window (B) (10-20 HPH) significantly restored backward movement to *unc-4;Ex[Punc-17::bPAC]* worms. Fisher’s Exact test vs *unc-4*, ** = p< 0.01, NS = not significant. N > 40 for each group. **C.** Ectopic expression of PDE-4/phosphodiesterase in VA motor neurons phenocopies the Unc-4 backward movement defect. (Left) The *Punc-4* promoter was used to drive expression of PDE-4/phosphodiesterase in VAs. Backward movement was scored in the tapping assay as either wild type (WT) or Uncoordinated (Unc). Worms expressing *Ex[Punc-4::PDE-4* displayed a strong backward Unc phenotype. Fisher’s Exact test vs wild type, **** = p< 0.0001. N>50. (Right) Representative tracks of ten *wild type* and ten *Punc-4::PDE-4* worms in a 3-minute period. Scale bar = 2mm. **D.** Ectopic expression of constitutively active GOA-1/GαO enhances the Unc-4 backward movement defect. The *Punc-4* promoter was used to drive expression of constitutively active GOA-1(Q205L) in VAs in *unc-4(ts)* worms. At 16°C, *unc-4(ts)* worms show wild type (WT) backward locomotion whereas *unc-4(ts)* worms that express GOA-1(Q205L) in VAs show uncoordinated (Unc) backward movement in the tapping assay. Fisher’s Exact test. *** = p< 0.001. N>50 for each genotype. **E.** GOA-1/GαO activation in VA motor neurons drives miswiring. Quantification of ectopic VA→AVB electrical synapses marked with UNC-7s::GFP. Data are percent of total VAs miswired with AVB electrical synapses (grey) vs. VAs that are not miswired (light blue) in *unc-4(e120), unc-4(e120); goa-1(lof), and unc-4(e120); goa-1(lof) ;Ex[Punc-4::GOA-1].* Note that loss of goa-1 function partially suppresses Unc-4 miswiring and that expression of wild-type GOA-1 (*Punc-4::GOA-1*) restores the AVB→VA gap junction defect. VAs included in analysis: VA2-4, VA8-10. Fisher’s exact test vs *unc-4(e120);goa-1(lof)*, *** = p<0.001.

### Reduced cAMP in VA neurons disrupts the backward movement circuit and induces miswiring with VA**→**AVB electrical synapses

Having established that forced elevation of cAMP is sufficient to rescue the backward movement defect of an *unc-4* mutant, we next asked if the reciprocal effect of reduced cAMP in VA neurons could induce the Unc-4 movement and miswiring phenotypes. To assess the effect of reduced cAMP on backward locomotion, we used the *unc-4* promoter to drive ectopic expression of the cAMP-specific phosphodiesterase, PDE-4, in VA motor neurons. In this experiment, worms expressing *Punc-4*::PDE-4 showed a strong Unc-4 like backward movement defect (**Figure 4C**) thus suggesting that reduced cAMP in VA neurons, which should arise from over-expression of PDE-4, is sufficient to impair backward locomotion. In a second experiment, we drove expression of the constitutively active allele of GOA-1(Q205L), which should antagonize cAMP synthesis (**Figure 3A**), in the VA neurons of a temperature-sensitive allele of *unc-4(ts)* (Miller et al., 1992). When grown at the permissive temperature (16°C), *unc-4(ts)* worms display wild-type backward movement. In contrast, expression of GOA-1(Q205L) in *unc-4(ts)* VA neurons resulted in a strong backward locomotion defect at 16°C (**Figure 4D**). Together, these experiments support the idea that lowered cAMP in VA neurons is sufficient to disrupt the backward movement circuit.

We performed an additional experiment as a direct test of whether reduced cAMP in VA neurons is sufficient to induce VA miswiring. In this case, we used the UNC-7S::GFP marker to confirm that the majority (83%) of VA neurons are miswired with VA→AVB gap junctions in the null allele, *unc-4(e120)* (**Figure 4E**). In addition, VA→AVB miswiring is substantially reduced (49%) in *unc-4(e120)* double mutants with *goa-1(lof)* (**Figure 5E**). Moreover, forced expression of wild-type GOA-1 in VA motor neurons (*Punc-4*::GOA-1) in *unc-4(e120); goa-1(lof)* mutants results in a VA→AVB miswiring defect (86%) comparable to that of *unc-4(120)* (**Figure 4E**). Together, these results are consistent with the hypothesis that cAMP acts cell autonomously within VA motor neurons to prevent miswiring with VA→AVB electrical synapses.

**Figure 5:**
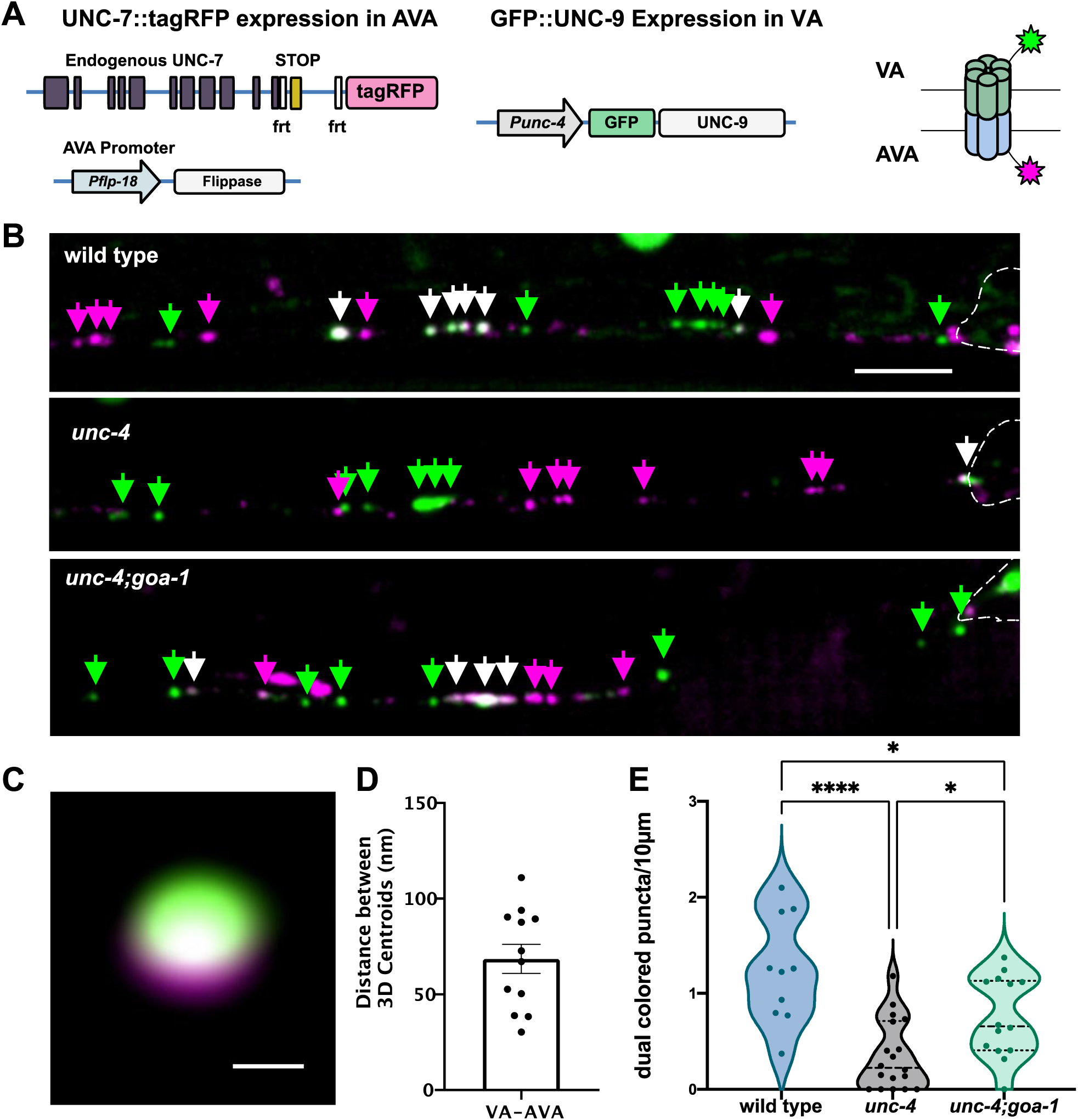
cAMP promotes neuron-specific assembly of electrical synapses. **A.** Two-color labeling of UNC-7 and UNC-9 innexins in heterotypic VA→AVA gap junctions. VA→AVA gap junctions contain UNC-7 (AVA) and UNC-9 (VA). AVA promoter (*Pflp-18*) drives flippase resulting in expression of endogenous UNC-7::tagRFP in AVA. VA promoter (*Punc-4*) drives expression of GFP::UNC-9 in VAs. Schematic of dual colored heterotypic gap junction. **B.** Representative images of AVA::UNC-7::tagRFP and VA::GFP::UNC-9 in VA process of wild-type, *unc-4(e2323)* and *unc-4(e2323);goa-1(lof)* in the ventral nerve cord of L4 stage larvae. Cell soma denoted by dashed outline. AVA::UNC-7::tagRFP (magenta arrowheads), GFP::UNC-9 (green arrowheads) and co-localized UNC-7 and UNC-9 (white arrowheads) puncta are denoted. Scale bar = 2.5 µm. **C.** Representative image of super resolution dual-colored gap junction. Scale bar = 200nm **D.** Quantification of 3D distance between centroids of UNC-7::tagRFP and GFP::UNC-9 images at dual colored puncta (see Methods). **E.** Quantification of density of dual-colored gap junctions. Violin plots for density of co-localized UNC-7::tagRFP and GFP::UNC-9 puncta in wild type (light blue), *unc-4(e2323)* (black) and *unc-4(e2323);goa-1(lof)* (green) VAs. Dashed line represents median. 2-way ANOVA. * = p < .05. **** = p <.0001. 561 has been pseudo-colored to magenta for all panels

### cAMP drives neuron-specific gap junction assembly

Our results show that cAMP antagonizes the formation of ectopic VA→AVB gap junctions (**Figure 3**). Because manipulations that elevate cAMP also restore backward locomotion to *unc-4* mutants (**Figure 4A**), we predicted that cAMP should also promote the formation of wild-type VA→AVA gap junctions that function in the backward movement circuit (**Figure 1**). To test this idea, we devised a live-cell imaging strategy to visualize VA→AVA gap junctions. VA→AVA gap junctions are composed of the innexins UNC-7 (expressed in AVA) and UNC-9 (expressed in VAs) (Starich et al., 2009). The resultant heterotypic gap junctions are composed of homomeric arrays of UNC-7 in AVA and UNC-9 in VA neurons. We utilized a two-color approach for live cell imaging of UNC-7 in AVA and UNC-9 in VAs. First, we expressed an N-terminal GFP fusion with UNC-9 in VA neurons (*Punc-4:*:GFP::UNC-9) (Meng et al., 2016). We observed that GFP::UNC-9 puncta largely reside in VA neuronal processes in the wild type but are preferentially displaced to VA soma in *unc-4* (**Figure S4**). These findings are consistent with previous EM reconstructions of wild-type and *unc-4* mutant VAs and thus suggest that VA::GFP::UNC-9 is a reliable marker of VA electrical synapses (White et al., 1986, 1992). Next, we used a flp/frt strategy for specific labeling of endogenous UNC-7 with TagRFP in AVA neurons (Schwartz and Jorgensen, 2016). With this two-color approach we could visualize both innexins, GFP::UNC-9 and UNC-7::TagRFP, that contribute to heterotypic VA→AVA gap junctions (**Figure 5A, B**).

We first used this strategy to monitor VA→AVA gap junctions in wild-type VAs. At the L4 stage, we observed multiple dual-color puncta in each VA (**Figure 5B, E**). Super resolution microscopy confirmed that GFP::UNC-9 in VA neurons and UNC-7::tagRFP in AVA are closely apposed (∼70nm) as expected for *bona fide* VA→AVA gap junctions (**Figure 5C**, **Video S2**) (See Methods) (Marsh et al., 2017; Oshima et al., 2013). Next, we tracked GFP::UNC-9/UNC-7::TagRFP puncta in *unc-4* mutant VAs and detected a significant decrease in the number of dual-color signal in the VA axon in comparison to wild type (**Figure 5B, E**). This observation confirmed previous results obtained from EM reconstruction showing that VA→AVA gap junctions are largely absent in *unc-4* mutants (White et al., 1992). Finally, we utilized our two-color labeling strategy to ask if elevated cAMP could restore VA→AVA gap junction assembly. For this test, we counted GFP::UNC-9/UNC-7::TagRFP puncta in *unc-4; goa-1(lof)* mutants and detected significant restoration of dual-color puncta in the VA axon in comparison to *unc-4.* Because the *goa-1(lof)* mutation is predicted to elevate cAMP, this observation supports the hypothesis that cAMP normally promotes VA→AVA gap junction biogenesis (Figure 3A).

### cAMP promotes the formation of functional neuron-specific VA→AVA electrical synapses

Functional gap junctions facilitate the rapid flow of ions between communicating neurons. Having determined that cAMP promotes the assembly of heterotypic VA→AVA electrical synapses composed of UNC-9 and UNC-7, we next asked if these VA→AVA gap junctions are functional. For this experiment, we devised an optogenetic strategy that uses the red-light sensitive channelrhodopsin, Chrimson, and Ca^2+^ sensor, GCaMP6s, to detect functional VA→AVA gap junctions. This approach relies on a previous finding that VA→AVA gap junctions are antidromic (Liu et al., 2017); ions flow unidirectionally from VA→AVA in opposition to the direction of cholinergic signaling from AVA to VA via chemical synapses. Thus, optogenetic excitation of VAs is predicted to activate AVAs via functional VA→AVA gap junctions and result in visible Ca^2+^ transients in AVA. To test this idea, we built a transgenic line expressing Chrimson in VA neurons (VA::Chrimson) and GCaMP6s in AVA (AVA::GCaMP6s) (**Figure 6A-B**). We determined that activation of VA::Chrimson in anteriorly placed VA neurons (VA2-4) triggers a robust GCaMP response in the AVA axon in the wild-type as predicted from previous electrophysiological recordings (**Figure 6C, Video S3**). This response was abrogated in *unc-7* mutants in which VA→AVA gap junctions are disabled as expected since UNC-7 is expressed in AVA and required for the formation of heterotypic gap junctions with UNC-9 in VA neurons (Starich et al., 2009). As an additional control, we determined that AVA GCaMP also depends on ATR, a necessary co-factor for optogenetic activation of Chrimson **(Figure S5**).

**Figure 6:**
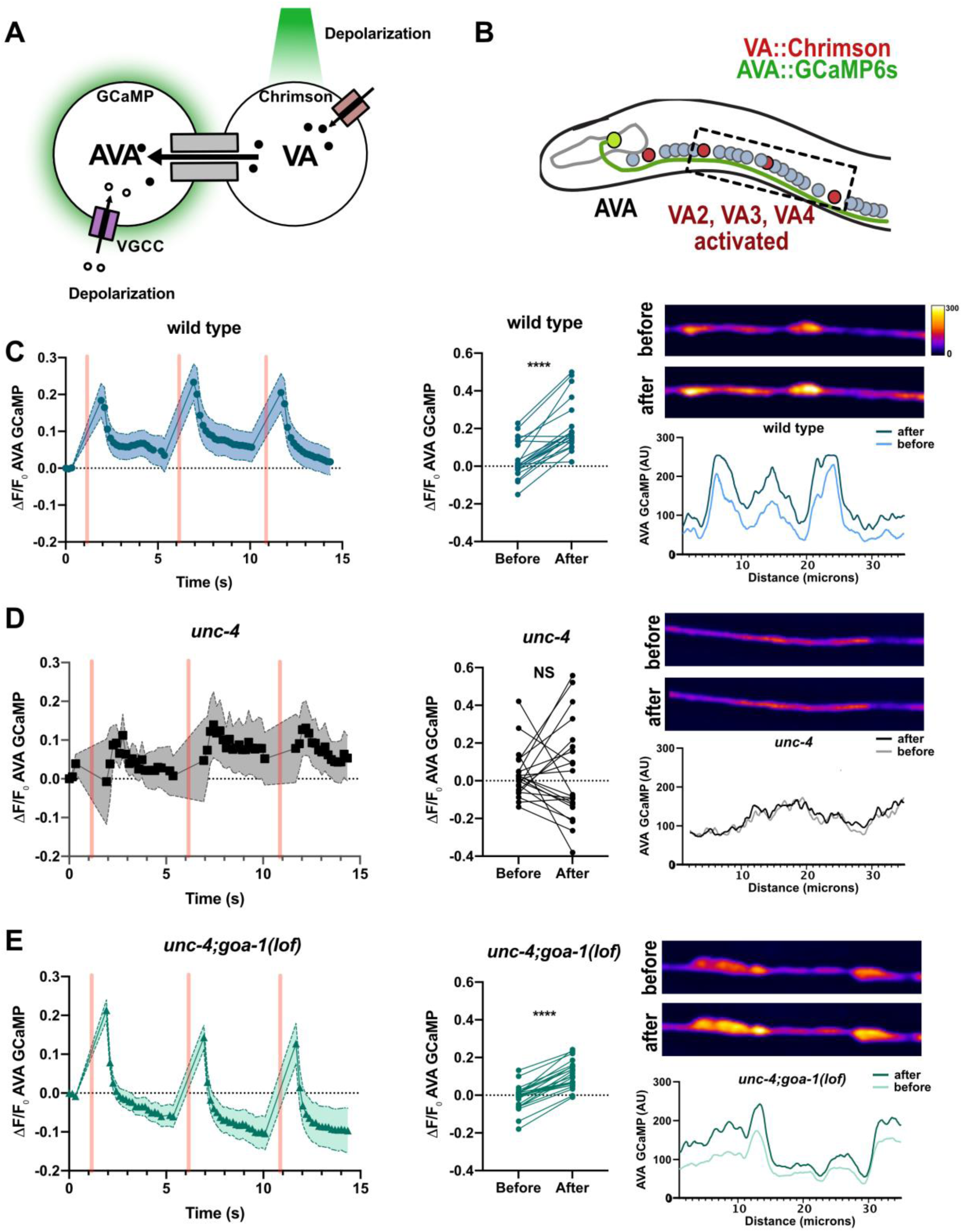
cAMP promotes assembly of functional VA→AVA electrical synapses. **A)** Schematic of strategy to monitor functional VA→AVA gap junctions. Red-shifted opsin, Chrimson (red rectangles) is expressed in VA motor neurons. Chrimson is activated over a wide spectral range from 550-650 (Basu et al., 2015; Klapoetke et al., 2014). The calcium sensor GCaMP6s is expressed in AVA. Exposure to green light (561 nm) activates Chrimson and depolarizes VAs leading to cation (black circles) flow through antidromic gap junctions with AVA (grey rectangles). The resultant AVA depolarization activates voltage gated calcium channels (VGCC) (purple rectangles) and calcium influx (open circles) which is detected by AVA-GCaMP. **B)** Schematic of assay for functional VA→AVA gap junctions. Each AVA neuron (green) located in the head extends an axonal process into the ventral nerve cord where it establishes antidromic (VA→AVA) electrical synapses with VA motor neurons. VA::Chrimson labels VAs (red) and AVA::GCaMP6s (green) marks AVA neurons. expression in VAs (*VA::Chrimson*) drives expression within VA motor neurons (red circles). A region posterior to the nerve ring containing VA2-VA4 (box) is activated with a 561nm laser and the GCaMP6s response in the adjacent AVA axon is measured with 488 nm excitation. **C)** Detection of wild-type VA→AVA electrical synapses. (**left**) Quantification of wild-type ΔF/F_0_ of *AVA::GCaMP6s* fluorescence over time. Three successive VA activations (500 ms) are denoted by pink vertical bars. Shaded area = SEM. N = 10 worms. (**middle**) Quantification of GCaMP6s ΔF/F_0_ before versus after 561 stimulation. N = 10 worms, 20 activations. Paired t-test. **** = p<.0001. (**Right**) Representative images of wild-type *AVA::GCaMP6s* signal before and after VA activation. Line scans of the representative images show *AVA::GCaMP6s* fluorescence intensity before and after VA activation. Heatmap indicates range of arbitrary units (AU) of fluorescence. **D)** VA→AVA electrical synapses are not detected in *unc-4(e2323).* (**left**) Quantification of *unc-4* AVA::GCaMP6s ΔF/F_0_ of *AVA::GCaMP6s* fluorescence over time. Three successive VA activations (500 ms) are denoted by pink vertical bars. Shaded area = SEM. N = 9 worms. (**middle**) Quantification of ΔF/F_0_ before versus after 561 stimulation. N = 9 worms, 18 activations. Paired t-test. NS = Not Significant. (**Right**) Representative images of *unc-4 AVA::GCaMP6s* signal before and after VA activation. Line scans of the representative images show *AVA::GCaMP6s* fluorescence intensity “before” and “after” VA activation. **E)** Functional VA→AVA electrical synapses are restored in *unc-4(e2323);goa-1(lof)* double mutants. (**left**) Quantification of *unc-4;goa-1* ΔF/F_0_ of *AVA::GCaMP6s* fluorescence over time. Three successive VA activations (500 ms) are indicated by vertical pink bars. Shaded area = SEM. N = 10 worms. (**middle**) Quantification of *AVA::GCaMP6s* ΔF/F_0_ before versus after 561 stimulation. N = 10 worms, 20 activations. Paired t-test. **** = p<.0001. (**Right**) Representative images of *unc-4;goa-1 AVA::GCaMP6s* expression before and after VA activation. Lines cans through the representative images show *AVA::GCaMP6s* fluorescence intensity before and after VA activation.

We next used this optogenetic assay to confirm that functional VA→AVA gap junctions are eliminated in *unc-4* mutants as predicted by EM reconstruction (White et al., 1992) and live-cell imaging (**Figure 5**). As expected, activation of VA::Chrimson in an *unc-4* mutant failed to trigger a detectable increase in AVA::GCaMP6s fluorescence (**Figure 6D**). We therefore concluded that our optogenetic assay can reliably detect functional VA→AVA gap junctions. Finally, we used this assay to ask: *Is cAMP sufficient to promote the formation of competent VA*→*AVA gap junctions?* Based on our behavioral (**Figure 3C**) and wiring results (**Figure 5B, E**), we predicted that elevating cAMP in an *unc-4* mutant should restore functional VA→AVA gap junctions. We assayed *unc-4; goa-1(lof)* double mutants and confirmed that activation of VA::Chrimson triggered a robust GCaMP response in AVA (**Figure 6E**). Taken together, these results argue that cAMP is sufficient to promote the assembly of functional VA→AVA electrical synapses.

### cAMP promotes gap junction trafficking in VA axons

In addition to the misassembly of VA electrical synapses with AVB, VA gap junctions are also misplaced to a different cellular compartment in *unc-4* mutants. In the wild-type animals, gap junctions with AVA are positioned on the VA axon whereas in *unc-4* mutants gap junctions with AVB are switched to the VA cell soma (**Figure 1A**) (Von Stetina et al., 2007; White et al., 1992). This change in the location of VA gap junctions led us to hypothesize that trafficking of gap junction subunits from the VA soma to the VA axon could be perturbed in *unc-4* mutants. To test this idea, we utilized GFP::UNC-9 to monitor gap junction trafficking in VAs. Time-lapse imaging revealed rapid bidirectional movement of GFP::UNC-9-labeled puncta in wild-type VA axons (**Figure 7A-B**). We observed both anterograde (average = 0.8 µm/sec) and retrograde movement (average = 1.3 µm/sec) with equal time in both directions (**Figure 7S**). In striking contrast, GFP::UNC-9 puncta were largely stationary in *unc-4* mutants and predominantly localized to the VA cell soma (**Figure 7A-B, Figure S6, Video S4**). To ask whether cAMP promotes gap junction trafficking, we monitored GFP::UNC-9 puncta in an *unc-4; goa-1(lof)* double mutant in which cAMP signaling should be elevated due to genetic ablation of GOA-1, a homeostatic antagonist of adenylate cyclase activity (**Figure 3A**). We observed a partial restoration of GFP::UNC-9 mobility in *unc-4; goa-1(lof)* mutants, a finding consistent with the hypothesis that cAMP promotes GFP::UNC-9 transport in VA neurons (**Figure 7A-D**).

**Figure 7:**
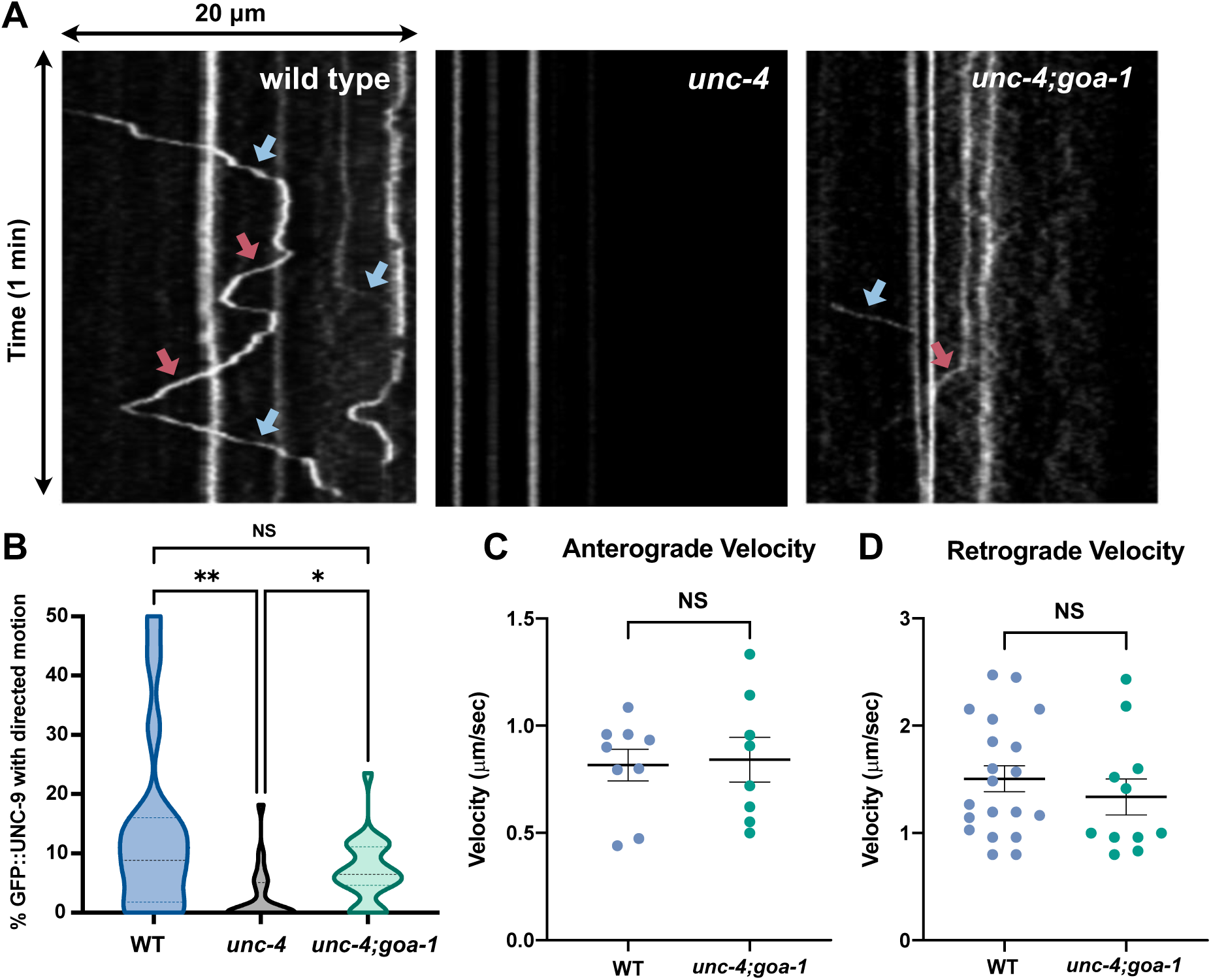
cAMP promotes axonal trafficking of UNC-9 gap junction components. **A)** *Punc-4::GFP::UNC-9* imaged over 3-minute period (200ms t-step) to monitor trafficking of UNC-9 in VA axonal process. Representative kymographs show retrograde (blue arrows) and anterograde (red arrows) trafficking of GFP::UNC-9 in wild type and in *unc-4(e2323); goa-1(lof)* worms. GFP::UNC-9 movement is substantially reduced in *unc-4.* **B)** Quantification of GFP::UNC-9 movement in wild-type (light blue), *unc-4(e2323)* (black) and *unc-4(e2323);goa-1(lof)* (green). Data are percent of motile puncta in a 3-minute period for given VA. N > 13 for each group. Kruskal-Wallis test. *** p= .0005, * p= .0326, NS= Not Significant. Results from VA2, VA3, VA4. **C)** Quantification of the anterograde velocity of individual GFP::UNC-9 puncta in wild-type (mean = 0.817 m/s) and *unc-4(e2323);goa-1(lof)* (mean = 0.842 µm/s) VA motor neurons. N=9 for each genotype. Student’s t-test, NS= not significant. **D)** Quantification of the retrograde velocity of individual GFP::UNC-9 puncta in wild-type (mean = 1.5 µm/s) and *unc-4(e2323);goa-1(lof)*(Mean = 1.34 µm/s) VA neurons. N=20 for wild type and N=10 for *unc-4;goa-1*. Student’s t-test, NS= not significant.

### UNC-4 gap junction assembly in VA axons during larval development

EM reconstruction, immunostaining experiments and live cell imaging have shown that the VA→AVA electrical synapses normally observed in the wild-type motor circuit are replaced with ectopic VA→AVB gap junctions in *unc-4* mutants (Schneider et al., 2012; Von Stetina et al., 2007; White et al., 1992) (**Figure 2H, Figure 5**). These results were obtained in late larval (L4) or adult worms, although VA motor neurons are generated at an earlier developmental stage (i.e., late L1 larvae). Notably, previous studies have determined that the adult pattern of connectivity in the motor circuit (e.g., VA→AVA gap junctions) is established by the L1-L2 larval molt (John White, personal communication, Sulston and Horvitz, 1977). Interestingly, experiments with a temperature sensitive *unc-4* mutant, suggest that UNC-4 function is required at a later stage, in L2-L3 larvae, to prevent VA miswiring (Miller et al., 1992). These observations suggested that UNC-4 might not be necessary for the initial assembly of VA→AVA gap junctions but rather could be required for their maintenance. We used our dual-color labeling strategy (**Figure 5A**) to confirm that VA→AVA gap junctions are detectable in L2 stage wild-type larvae (**Figure 8A, B**). Interestingly, we observed a comparable number of VA→AVA electrical synapses at the L2 stage in *unc-4* mutants. This finding is consistent with the hypothesis that UNC-4 is not required for the initial formation of VA→AVA gap junctions. In the wild type, a substantial number of additional VA→AVA gap junctions are detectable later in development by the L4 stage (**Figure 8B**). However, we observed no comparable increase in VA→AVA gap junctions in *unc-4* mutants (**Figure 8B**). Together, these findings suggest that UNC-4 promotes the formation of additional VA→AVA gap junctions during larval development as VA neurons expand in size.

**Figure 8:**
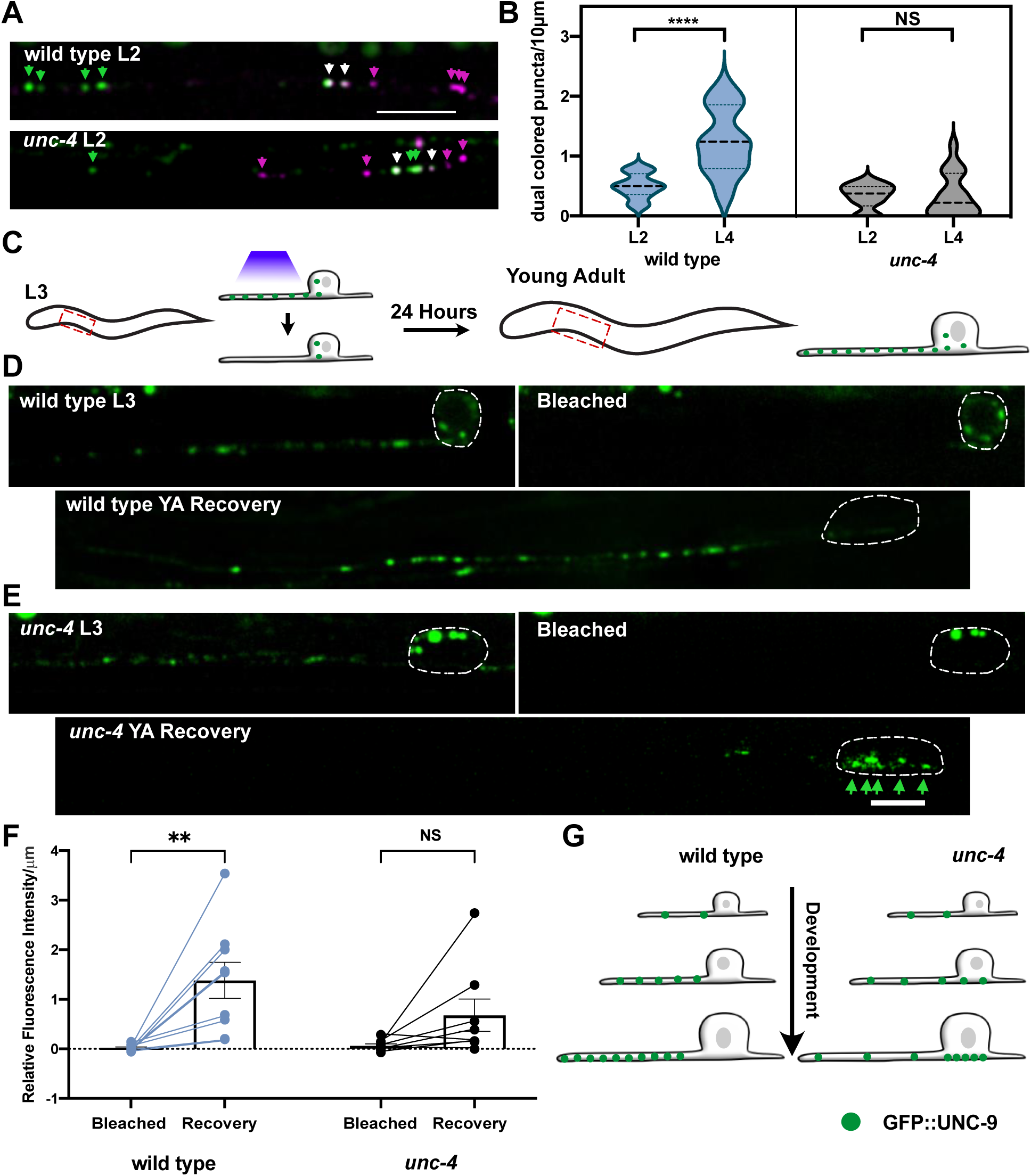
UNC-4 promotes gap junction assembly in VA axon during larval development. **A)** Representative images of AVA::UNC-7::tagRFP and VA::GFP::UNC-9 in VA axons of wild type and *unc-4* in L2 larvae. AVA::UNC-7::tagRFP (magenta arrowheads), GFP::UNC-9 (green arrowheads) and co-localized UNC-7 and UNC-9 (white arrowheads) puncta are denoted. Scale bar = 2.5 µm. **B)** Quantification of dual-colored gap junctions throughout development. Violin plots for density of co-localized UNC-7::tagRFP and GFP::UNC-9 puncta in wild-type (light blue) and *unc-4(e2323)* (black) VAs in L2 and L4 larvae. Wild-type, but not *unc-4*, VAs have significantly more dual-colored puncta at L4 compared to L2. L4 data for wild type and *unc-4(e2323)* also depicted in Figure 5E. Dashed line represents median. 2-way ANOVA, **** = p <.0001. NS = Not Significant. **C)** Schematic of FRAP strategy. Individual L3 larvae are immobilized and a single VA axon (VA2, VA3, or VA4) is bleached. Each treated animal is recovered for growth on NGM plate until young adult (24 hour) and then re-imaged to record fluorescence intensity from photobleached VA axon. Representative images of VA::GFP::UNC-9 puncta in **D)** wild type and **E)** *unc-4(e2323)* at L3 before and after photobleaching and at the young adult. Dashed outline denotes VA soma. Scale bar = 2.5 µm. Green arrowheads mark accumulated GFP::UNC-9 on soma. **F)** Quantification of relative fluorescence intensity of individual VAs immediately after photobleaching (Bleach) and after 24 hours (Recovery) in wild type and *unc-4(e2323).* Plots of intensity are normalized to VA axon length in each case. Bleach and Recovery represented as their fluorescence intensity value relative to fluorescence intensity prior to bleaching. 2-way ANOVA, ** p = .0023. NS = Not Significant. **G)** Schematic of VA motor neuron during larval development. Additional GFP::UNC-9 puncta accumulate in wild-type VA process during larval development. In *unc-4* mutant VA neurons, GFP::UNC-9 puncta accumulate on the cell soma during larval development.

Having observed that UNC-9 trafficking is drastically curtailed in *unc-4* mutant VA neurons at the L4 stage (**Figure 7**), we hypothesized that the failure to assemble additional VA→AVA gap junctions in *unc-4* mutants could be due to defective gap junction trafficking during development. To test this idea, we performed a Fluorescence Recovery After Photobleaching (FRAP) experiment. If gap junction components are actively trafficked into wild-type VA motor axons during development, then GFP::UNC-9 fluorescence should recover after an early photobleaching event. For this experiment, we tracked GFP::UNC-9 at three timepoints: 1) in L3 larvae prior to photobleaching; 2) in L3 larvae immediately following photobleaching; and 3) in young adults, 24- hour after photobleaching.(**Figure 8C**). This experiment revealed that the GFP::UNC-9 signal was fully restored in wild-type VAs in young adults. In fact, average GFP::UNC-9 fluorescence actually exceeded initial values before bleaching in the L3 which we attribute to enhanced trafficking of GFP::UNC-9 during development (**Figure 8D, F**). In *unc-4* mutants, however, we observed no significant recovery of fluorescence following photobleaching (**Figure 8E, F**), thus indicating that little GFP::UNC-9 is transported into *unc-4* mutant VA axons during this period. Together, these results suggest that UNC-4 preserves the specificity of VA electrical synapses by promoting transport of UNC-9 into VA axons for the formation of additional VA→AVA gap junctions during larval development (**Figure 8G**).

## Discussion

Electrical synapses, or gap junctions, are fundamentally important to neural function, but little is known of the mechanisms that regulate the neuron specificity or subcellular placement of gap junctions. Here, we demonstrate that cAMP directs both the specificity and placement of electrical synapses in the *C. elegans* motor circuit. Because experiments in cultured mammalian cells have also identified cAMP as a regulator of gap junction biogenesis (Atkinson et al., 1986; Holm et al., 1999; Paulson et al., 2000; Solan and Lampe, 2016; TenBroek et al., 2001; Thévenin et al., 2017), we suggest that cAMP directs evolutionarily conserved pathways that control the assembly of electrical synapses in the brain (**Figure 9**).

**Figure 9:**
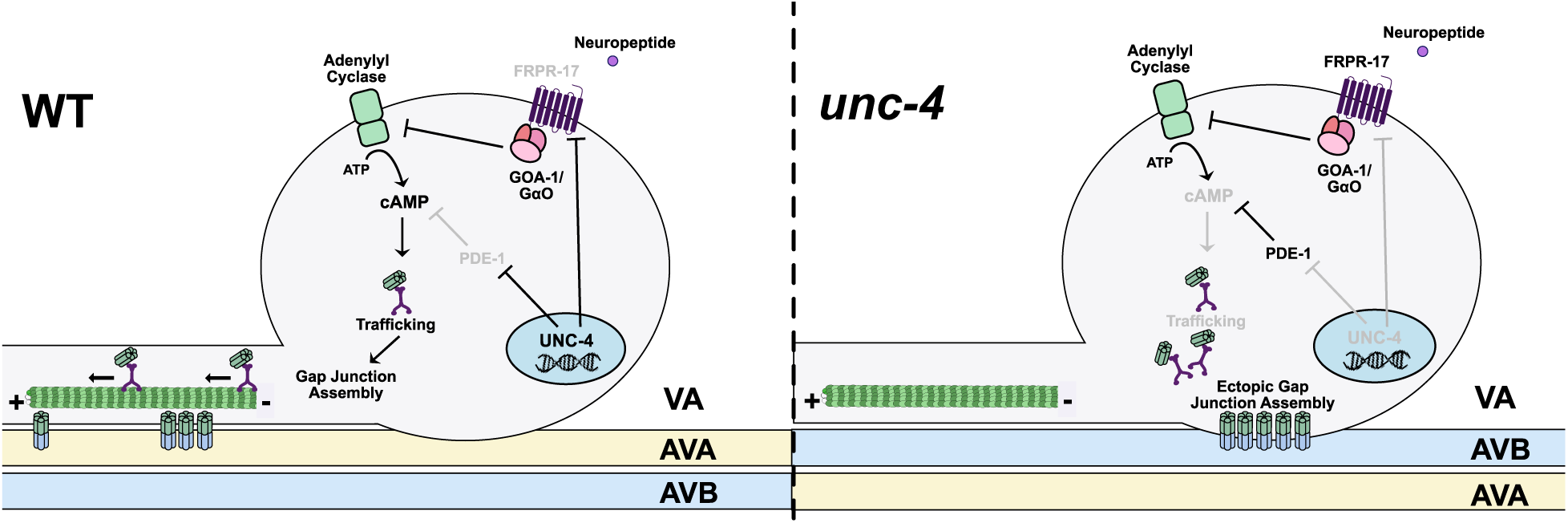
UNC-4 regulates cAMP to direct trafficking, neuron-specific assembly and subcellular placement of electrical synapses. (**Left**) In wild-type VAs, UNC-4 blocks expression of cAMP antagonists, PDE-1 (phosphodiesterase) and FRPR-17(GOA-1-coupled GPCR), to maintain cAMP. cAMP promotes trafficking of UNC-9 into the VA axon for assembly with UNC-7 at heterotypic VA→AVA electrical synapses during larval development. (**Right**) Ectopic expression of PDE-1 and FRPR-17 in *unc-4* mutant VAs reduces cAMP levels and limits UNC-9 trafficking. UNC-9 accumulates in VA soma for assembly with UNC-7 at heterotypic VA→AVB gap junctions.

### The neuron-specificity of electrical synapses is tightly regulated

The formation of functional electrical synapses depends on expression of compatible gap junction subunits (e.g., connexins/innexins) in each of the coupled neurons. Although necessary for gap junction assembly, connexin/innexin expression is not sufficient to explain the striking neuron specificity of electrical synapses (Martin et al., 2020) For example, in the mouse retina, photoreceptor and bipolar neurons are closely apposed but are not electrically coupled despite the joint expression of connexin Cx36 for gap junction assembly with other adjacent neurons (Asteriti et al., 2017; Deans et al., 2002; Trenholm and Awatramani, 1995). Similar examples of selective gap junction assembly among neurons that express compatible connexins have been observed in the carp retina and mouse neocortex (Greb et al., 2017; Yao et al., 2016). To investigate this question in *C. elegans*, we relied on uniquely available comprehensive catalogs of neuron-specific synapses and innexin expression for the entire nervous system (Bhattacharya et al., 2019; Cook et al., 2019; Taylor et al., 2020; White et al., 1986). We used the wiring diagram to identify all pairs of neurons with chemical synapses but not gap junctions. We then determined the subset of neuron pairs that express the compatible innexins, UNC-7 and UNC-9 (Starich et al., 2009). This straightforward approach identified over 600 pairs of adjacent neurons that create chemical synapses but fail to form gap junctions despite their close proximity and co-expression of UNC-7 and UNC-9 **(Supplemental Table 2**). Together, these observations argue that the neuron specificity of electrical synapses across species must depend on regulatory pathways that control gap junction assembly.

Recent studies *in vivo* have identified molecular determinants of gap junction biogenesis. In the zebrafish, the connexin-associated protein, Zona Occludens 1 (ZO1), localizes to sites of gap junction formation in the Mauthner neuron where it is proposed to function as a scaffolding protein for gap junction assembly (Lasseigne et al., 2021; Marsh et al., 2017). In *C. elegans*, the membrane protein, NLR-1/CASPR, is similarly proposed to recruit innexins to sites of gap junction assembly in multiple tissues including neurons. It is unclear, however, if either ZO-1 or NLR-1/CASPR are integral components of all gap junctions or are uniquely required for directing the assembly of gap junctions with between specific cell types.

### cAMP directs the neuron specificity of electrical synapses

In neurons, cAMP regulates morphological polarity, axon guidance and synaptic function (Hannila and Filbin, 2008; Lee, 2015). Both cAMP synthesis and degradation are actively modulated to tune cAMP-dependent signaling. G-protein-coupled receptors (GPCRs) influence cAMP production by either promoting (GαS) or inhibiting (Gαi/o) adenylyl cyclase-dependent conversion of ATP to cAMP (**Figure 3A**). Phosphodiesterases (PDEs) down regulate cAMP signaling by cleaving cyclic phosphodiester bonds to convert cAMP to AMP. Our results show that the UNC-4 transcription factor represses antagonists of cAMP signaling, a phosphodiesterase (PDE-1) and a GPCR (FRPR-17), in VA motor neurons to maintain neuron-specific electrical synapses required for wild-type movement. In *unc-4* mutants, when PDE-1 and FRPR-17 are ectopically expressed, specific VA neurons fail to connect with AVA and instead form electrical synapses with AVB interneurons (VA→AVB) (**Figure 1**). This effect is reversed by mutations that disable either *pde-1* or *frpr-17* in an *unc-4* mutant background, a finding that directly implicates *pde-1* and *frpr-17* in VA miswiring (**Figure 2**). Similar suppression was obtained from a series of experiments that elevate cAMP signaling, i.e., treatment with the cAMP analog, 8-Br-cAMP (**Figure 3B**) optogenetic activation of adenylyl cyclase (**Figure 4**), loss-of-function mutation in GαO and gain-of-function mutations in either adenylyl cyclase or GαS (**Figure 3F**). Overall, our findings are consistent with the hypothesis that cAMP signaling normally favors the wild-type pattern of electrical synapses in VA motor neurons (**Figure 3**).

Although our studies focused on a subset of VA neurons in the anterior ventral nerve cord (VA2, VA3, VA4), cAMP signaling may be broadly required to block the formation of dysfunctional VA→AVB gap junctions. We observed, for example, that a loss-of-function mutation in GOA-1/ GαO prevents the formation of VA→AVB gap junctions in VA2-VA4 as well as in additional posteriorly located VA neurons in *unc-4* mutants (data not shown). The fact that ectopic expression of the proposed GOA-1-coupled GPCR, FRPR-17, is limited to VA2 in *unc-4* mutants (**Figure 2**), suggests that additional GPCRs are likely to function in posteriorly located VAs to promote GOA-1 activity. Because FRPR-17 is predicted to bind FMRFamide ligands, we suggest the intriguing possibility that neuropeptide signaling may regulate the neuron specificity of electrical synapses in the VA motor circuit.

Our results suggest that cAMP in VA motor neurons is modulated by antagonistic G-protein signaling pathways to control electrical synaptic specificity. GSA-1/GαS promotes wild-type VA→AVA electrical synapse formation, whereas GOA-1/Gαi/o favors ectopic VA→AVB gap junction assembly. These antagonistic pathways are reminiscent of a previously described program that regulates meiotic diapause in *C. elegans*. During ovulation, gap junctions between sheath cells and oocytes are destabilized to enable oocyte maturation. This effect is regulated by opposing G-proteins which are proposed to either promote (GSA-1/GαS) or antagonize (GOA-1/Gαi/o) gap junction destabilization (Govindan et al., 2006, 2009). Thus, the antagonistic action of G-protein signaling mechanisms could be a consistent motif for modulating cAMP control of gap junction assembly and function.

### Subcellular placement and maintenance of electrical synapses

The placement of electrical synapses in neural circuits must depend on the shipment of gap junction components to the site of assembly but the mechanisms that regulate gap junction trafficking in the nervous system are largely unknown. Studies in cultured cells have established that connexins are exported from the Golgi apparatus in vesicles which are then transported along microtubules for assembly at gap junction plaques. Kinesin motors likely drive microtubule transport of connexin-containing vesicles although evidence for this role is indirect (Flores et al., 2012; Fort et al., 2011; Shaw et al., 2007). Additional studies *in vitro* have reported cAMP-dependent expansion of gap junction plaques which could result from enhanced trafficking of gap junction components (Paulson et al., 2000; TenBroek et al., 2001). Our findings in the *C. elegans* motor circuit parallel these observations and suggest that cAMP signaling promotes assembly of electrical synapses in the nervous system by regulating gap junction trafficking.

The specific subcellular location of an electrical synapse is linked to its role in neuronal function. For example, the axon initiation segment is densely populated with voltage gated ion channels and gap junctions in this region can amplify local action potentials, a property important for fast network oscillations (Connors, 2017; Schmitz et al., 2001; Simon et al., 2014; Thomas et al., 2020; Traub et al., 2002). Dendro-dendritic gap junctions can be similarly positioned near voltage-gated ion channels for synergistic interactions that amplify currents (Connors, 2017; Sotelo et al., 1986; Zsiros et al., 2007). In some cases, specific connexins are targeted to different subcellular compartments for gap junction assembly. For example, in the zebrafish Mauthner neuron, Cx35.5 is directed to the axon whereas Cx34.1 is positioned in a lateral dendrite for each to establish electrical synapses with different coupled neurons (Miller et al., 2015, 2017). Our results show that cAMP is required for active transport of the innexin UNC-9 from the VA neuron cell soma into the axon where it is assembled into VA→AVA electrical synapses. In contrast, at low levels of cAMP, trafficking is curtailed and UNC-9 remains in the cell soma where it establishes gap junctions with the AVB interneuron. Important unanswered questions include the cell biological mechanism of cAMP-dependent trafficking and whether additional factors, potentially regulated by UNC-4, are required for the neuron specificity of UNC-9 gap junction assembly in the VA axon (AVA) vs cell soma (AVB).

Studies of developing neural circuits have determined that initial patterns of connectivity are maintained with the addition of new synapses as neurons expand in size. Morphological analysis of a developing nociceptive circuit in Drosophila revealed that the number of chemical synapses among established synaptic partners increases with larval growth. (Gerhard et al., 2017). Similarly, existing connections in the *C. elegans* nervous system are strengthened during larval development with the addition of new synapses that scale in proportion to increasing neurite length (Witvliet et al., 2020). Our findings point to a similar phenomenon for electrical synapses. In the wild type, in the number and density of VA→AVA gap junctions increase during larval development. Interestingly, a few VA→AVA gap junctions are initially established in *unc-4* mutants in early L2 larvae, but these fail to expand with larval growth, likely due to defective trafficking (**Figure 8**). This finding is consistent with the previous observation that UNC-4 function is required after the initial establishment of the adult pattern of connectivity in the ventral cord motor circuit in late L1 larvae (Miller et al., 1992, J. White, personal communication). Together, these results suggest that UNC-4 controls a genetic program that promotes insertion of additional neuron-specific gap junctions in the VA neurite to sustain functional electrical connectivity as the motor circuit expands in size.

In conclusion, we have used live-cell imaging, an *in vivo* assay of electrical synaptic function and genetic analysis to establish that cAMP controls a trafficking mechanism that directs the subcellular placement and neuron specificity of electrical synapses in the developing *C. elegans* motor circuit. The widely reported role of cAMP signaling in gap junction assembly and function in vertebrate cells suggests that cAMP-dependent pathways are also likely to govern the formation and maintenance of electrical synapses in mammalian neural circuits.

## Acknowledgments

We thank O. Hobert, A.Gottschalk and D. Yan for reagents and Eduard Tataru for his help. This work was funded by NIH grant R01NS113559 to DMM. SP received fellowship support from American Heart Association (19PRE34380582) and the National Science Foundation (DGE:1445197). Flow Cytometry experiments were performed in the Vanderbilt Flow Cytometry Shared Resource which is supported by the Vanderbilt Ingram Cancer Center (P30 CA68485) and the Vanderbilt Digestive Disease Research Center (DK058404). Some strains were provided by the CGC, which is funded by NIH Office of Research Infrastructure Programs (P40 OD010440).

## Author Contributions

Conception and design of experiments: S.P., D.M.M.; Generation of transgenic lines: S.P., R.S., S.VS.; Collected fixed images and quantified results: S.P., S.H., R.S.; Performed live imaging and quantified results: S.P. Sorted cells for RNA-Seq: R.M.; Performed RNA-Seq Analysis: S.P.; Screened for *unc-4* suppressors: S.P., I.S., A.M.; Wrote the final document: S.P., D.M.M. Critically revised manuscript and approved final version for publication: S.P., R.S., S.VS., A.M., I.S., S.H., R.M., D.M.M.

## STAR METHODS

### RESOURCE AVAILABILITY

#### Lead contact

Further information and requests for reagents should be directed to and will be fulfilled by the lead contact, David M. Miller III

#### Materials availability

Key *C. elegans* strains in this study have been deposited to the CGC.

#### Data and code availability

The RNA-seq datasets generated during this study are available at GEO (GSE173287)

### EXPERIMENTAL MODEL AND SUBJECT DETAILS

#### Strains and Genetics

*C. elegans* strains were grown at 23° C unless otherwise noted on OP50-1 *Escherichia coli*- seeded nematode growth medium plates (Brenner, 1974). For RNAseq experiments, *C. elegans* strains were grown on 8P nutrient agar seeded with *E. coli* strain NA22. The N2 strain was used as the wild-type reference. *unc-4(e2323)* was used for all experiments containing *unc-4* unless otherwise noted in the text. Mutant alleles and strains used in this study are described in the Key Resources Table.

### METHOD DETAILS

#### Molecular Biology

We used the In-Fusion cloning kit (Takara) to build all transgenes in this study (**Key Resources Table**). Plasmids were injected into N2 lines before crossing into a given genotype. The chromosomal integrants, *wdIs117(pUnc-4::Chrimson), ufis26[pUnc-4::mCherry],* and *wdIs90[pUnc-4C::GFP]* were obtained by x-ray irradiation (Miller and Niemeyer, 1995)and outcrossed for three generations.

#### Bulk RNA sequencing of FACS-isolated cells

We used Fluorescence Activated Cell Sorting (FACS) to isolate VA neurons labeled by an intersectional strategy. An integrated strain expressing *ufis26[pUnc-4::mCherry]* (VAs, DAs, SAB, I5, AVF) and *wdIs90[pUnc-4C::GFP]* (DAs, SAB, I5) labeled VA and AVF exclusively with mCherry in wild-type and *unc-4* L2 larvae (NC2957, NC2958). With 12 VA neurons (VA1-VA12) vs 2 AVFs (AVFL, AVFR) in each animal, we reasoned that VAs would be enriched relative to AVF with this strategy. Cell dissociation and FACS were performed as previously described (Spencer et al., 2014; Taylor et al., 2020). Briefly, synchronized L1 larvae were plated on 8P plates seeded with NA22 and grown overnight at 23 C. L2 larvae were then dissociated by successive treatments with solutions of 0.25% SDS, 0.2M DTT and 15mg/mL pronase and passed through a 5µm filter to remove debris before FACS. Dead cells were excluded by DAPI staining. VA and AVF neurons (mCherry plus, GFP minus) were collected in Trizol for RNA extraction. Three biological replicates for each genotype were performed with >50,000 cells/sample. For the wild-type VA expression profile, a reference sample of all cells was obtained from quick frozen aliquots of synchronized L2 larvae. The Clontech-Takara SMART-Seq V3 Ultra Low Input RNA Kit was used for cDNA synthesis and amplification. Paired-end-100 (PE-100) data were collected in an Illumina NovaSeq 6000. ≥ 50 million reads/sample were obtained for three independently-isolated VA samples from both wild-type and *unc-4* larvae and for three L2 whole animal reference samples. Reads were analyzed using CLC Genomics Workbench Version 11. Differentially-expressed transcripts were obtained using the RNA-Seq Differential Expression analysis pipeline in CLC, which utilizes a negative binomal GLM model. Genes were considered significantly regulated if expression was >2-fold and FDR-p-value <.01 compared to control.

#### Feeding RNA Interference Experiments

Bacteria producing double-stranded RNA for each target gene (**Table S1**) were seeded on NGM plates. Adult *unc-4(e2323);eri-1;lin-15* worms were plated on the seeded plates and allowed to lay eggs for two hours and progeny were grown at 23°C. L4 progeny were picked to a new plate for backward movement assays (see below). Experimenters were blinded to genotype. Either *goa-1(lof)* or *ceh-12(lof)* were used in each experiment as positive controls for RNAi-dependent suppression of the Unc-4 backward movement defect.

#### Pharmacological elevation of cAMP signaling

NGM plates were seeded with OP-50 containing 0.5mM 8-Bromoadenosine 3′,5′-cyclic monophosphate (8-Br-cAMP) (Sigma Aldrich) (Hussey et al., 2017). Plates were kept in the dark to prevent degradation of 8-Br-cAMP and used within one week. *unc-4(e2323)* adults were allowed to lay eggs on 8-Br-cAMP-containing plates for 2-hours and then removed. Progeny were grown to the L4 stage on 8-Br-cAMP-containing plates and transferred to NGM-plates seeded with OP-50 for behavioral analysis. Controls were *unc-4(e2323)* L4 larvae grown on NGM-plates seeded with OP-50. The experimentalist was blinded to growth condition.

#### Optogenetic activation of cAMP

NC3815 worms (*unc-4(e2323);lite-1(ce314*); zxIs53[*punc-17::bPAC::YFP, pmyo-2::mCherry*]) were grown on ATR-containing NGM-plates (Steuer Costa et al., 2017) were allowed to lay eggs for 2 hours to produce tightly synchronized larvae. Larvae were subjected to blue light activation (A) 0-10 HPH (Hours Post Hatch), (B) 10-20 HPH, and (C) 20-30 HPH. Larvae were maintained in a darkened room for a series of pulses of blue light (70 µW/mm²) of 10s on followed by 10s off for the 10 hr duration of each treatment period (Steuer Costa et al., 2017). Following stimulation, L4 worms were transferred to a new NGM plate for a tapping assay performed at the young adult stage (see below) for backward movement. NC3815 kept in the dark were used as negative controls. The experimentalist was blinded to genotype. Fisher’s exact test was used to determine significance.

#### Behavioral Assays

Backward movement was assessed by either 1) a “tapping assay” or 2) video tracking software (WormLab).

##### Tapping Assay to assay backward locomotion

L4 larvae were tapped on the head with a platinum wire to evoke backward locomotion (Von Stetina et al., 2007). *unc-4* mutants typically coil dorsally with head tap (White et al., 1992) and are unable to sustain backward movement. Movement was scored as “wild-type” for sustained sinusoidal backward locomotion (two full bends) or “Unc” for failed backward locomotion. Experimenters were blinded to genotype. N> 50 for each genotype. Fisher’s exact test was used to determine significance. (**Figures 2B, 4B-D**).

##### Video Tracking

L4 larvae were plated on a lightly-seeded plate of NGM with OP-50. WormLab was used to track the plated worms over a three-minute period with a 1s blue light pulse at 5s intervals throughout to promote movement. *unc-4* mutants exhibit a readily detectable reduction in backward distance traveled in a three-minute period compared to wild-type (Figure 1). We therefore used this to assay to identify suppression of the Unc-4 backward movement defect. Worms were included in analysis if they were captured and tracked for the entire 3-minute period of the video. A 2-way-ANOVA was used to determine significance. (**Figures 1C, 3B, 3E**).

#### Microscopy and Image Analysis

##### Immunostaining to detect AVB gap junctions with ventral cord motor neurons

AVB gap junctions in the ventral nerve cord were marked with *wdIs54[Punc7::UNC-7S::GFP* and immunostained to detect the dim GFP signal (Starich et al., 2009; Von Stetina et al., 2007) as previously described (Finney and Ruvkun, 1990). Briefly, L4 larvae were successively immersed in sealed microfuge tubes in liquid nitrogen for three freeze-thaw cycles before incubating in 1%paraformaldehyde for 40 minutes. Following fixation, a series of treatments (1% BME, 10 mM DTT, 0.3% H_2_O_2_, 0.1% Triton-X) were performed to permeabilize the cuticle. Treated larvae were incubated with 1:500 anti-GFP primary antibody (Roche, 11814460001) overnight at 4°C. Following washes with Antibody Buffer B (AbB), larvae were incubated with 1:500 goat-anti-mouse-Cy3 (Jackson ImmunoLaboratories, AB_2338680) for 2-hours at room temperature, washed with AbB, stained with DAPI (1:1000) for 30 minutes and mounted with VectaShield (Vector Labs). Z-stack (0.2 µm steps) images of DAPI (405 nm excitation) and Cy3 (561 nm excitation) were obtained in a Nikon A1R confocal microscope with a 60X objective (Plan Apo Lambda Oil, 1.40=NA). Images were 3D-deconvolved using Nikon NIS elements. VA neurons were identified by position in the DAPI-stained queue of ventral cord nuclei (Miller and Niemeyer, 1995). Cy3-stained puncta adjacent to VA nuclei were scored as gap junctions with AVB. To ensure that puncta were not background, we imposed a size and fluorescence intensity cutoff using NIS Elements. N> 20 VA neurons were scored for each group. Fisher’s exact test was used to determine significance.

##### Single molecule mRNA Fluorescence In Situ Hybridization (smFISH)

smFISH was performed with custom *pde-1* and *frpr-17* probes linked to Quasar® 670 or 561 respectively (Biosearch Technologies). Synchronized L2 larvae were collected by washing plates with M9, fixed in 4% paraformaldehyde in 1X PBS for 45 min and permeabilized in 70% ethanol for 24-48 h. Hybridization followed the manufacturer’s instructions (http://www.biosearchtech.com/stellarisprotocols) and was performed at 37°C for 16h in Stellaris RNA FISH hybridization buffer (Biosearch Technologies Cat# SMF-HB1-10) containing *pde-1 or frpr-17* probe at 1:100. VA neurons were marked with *Pbnc-1::GFP* (OH15624). Cell nuclei were stained with DAPI. Z-stacks were collected in a Nikon spinning disk confocal microscope equipped with optical filters for DAPI, Quasar® 670 or 651 and GFP using a 100X objective (NA=1.49) in 0.2 μm steps spanning the cell body and merged for quantification following 3D- deconvolution in NIS elements. smFISH puncta were counted if they corresponded to circular fluorescent spots, exceeded the Quasar® 670/561 background signal and were located within a GFP-labeled VA cell body. At least 20 worms were scored for each group and the Mann-Whitney test used to determine significance.

##### Dual-colored heterotypic VA→AVA gap junction

Worm lines expressing endogenous AVA::UNC-7::tagRFP and transgenic VA::GFP::UNC-9 were imaged to identify VA→AVA gap junctions. To build these lines, UNC-7 was endogenously tagged at the N-terminus with frt-STOP-UTR-frt-tagRFP (*syb2341*). The *Pflp-18* promoter was used to drive filppase in AVA resulting in tagged UNC-7::tagRFP selectively in AVA. C-terminal-GFP-fused UNC-9 (Meng et al., 2016) was expressed in VAs using the UNC-4 promoter (*Punc-4::GFP::UNC-9*). L4 worms were placed on 10% agarose pads and immobilized with 50mM muscimol. Z-stacks were captured (0.2 µm/step) for the width of each VA process (VA2, VA3, VA4) for GFP (488 nm excitation) and tagRFP (561 nm excitation) Stacks were 3D-deconvolved in NIS elements. Images were thresholded by fluorescence intensity, circularity, and size to identify puncta in each color in NIS elements. Puncta were counted as dual-colored if puncta of both colors were at the same point. The number of dual-colored puncta was normalized by distance for each VA counted to yield a density value (# puncta/10 μm) (density). A 2-way ANOVA was used to determine significance.

##### Structured Illumination Microscopy

Animals fixed with 4% paraformaldehyde for thirty minutes (NC3775 [unc-7(syb2341);*Ex[Pflp-19::flppase, Punc-4::GFP::UNC-9]*) were mounted with Vectashield and 1.5 coverslips. Z-stacks (0.07 µm/step) of GFP (488 nm excitation) and tagRFP (561 nm excitation) were captured with Nikon N-SIM microscope 100X SR Apo TIRF (1.49 NA) objective in 3D-SIM mode. Slice reconstruction was performed using NIS Elements. Following reconstruction, 3D mask for each fluorophore was created based on intensity. The centroid of each 3D mask was determined and the distance between the centroids was measured to estimate the proximity of UNC-7 vs UNC-9 gap junction arrays.

##### Monitoring functional VA→AVA gap junctions

We constructed a transgenic line (NC3666 [wdIs117(*Punc-4*::Chrimson);Ex1148[*pFlp-18::GCaMP6S*]) in which *Punc-4* drives expression of the red-light-activated opsin Chrimson in VAs and the *Pflp-18* promoter drives expression of the Ca^2+^ sensor GCaMP6s in AVA. Worms were grown on plates containing All-trans-Retinol (ATR), a necessary cofactor for Chrimson unless otherwise noted. L4 worms were mounted on 10% agarose pads and anesthetized with 50mM muscimol and a slurry of 0.5 µm Polybead® Carboxylate Microspheres (Polysciences). We captured AVA::GCaMP at 10 frames/sec (100ms) with a 100X SR Apo TIRF (1.49 NA) objective on a Nikon Spinning Disk. VA::Chrimson was activated with a 500 ms burst of 561 nm every 5 seconds. Following Chrimson activation, the next AVA::GCaMP frame was captured within 500 ms, Importantly, t^1/2^ = 600ms for GCaMP6s (Chen et al., 2013). AVA::GCaMP signal from the AVA process in a region adjacent to VA2-VA4. A single Z-plane was captured for 1-minute. Movies were 2D deconvolved and aligned using NIS elements. AVA fluorescence intensity was corrected by subtracting the value of a background ROI adjacent to the AVA process of the same size for each time point. Change in fluorescence intensity was calculated as ΔF/F_0_ = (F_t_-F_0_)/F_0_ (F_0_ = baseline fluorescence intensity of timepoint three frames before first Chrimson activation. F_t_ = Fluorescence intensity of a given timepoint). To detect evoked changes in AVA::GCaMP, we compared the ΔF/F_0_ of AVA::GCaMP immediately before Chrimson activation versus the ΔF/F_0_ of AVA::GCaMP immediately following Chrimson activation A paired t-test was performed for AVA GCaMP fluorescence time points before vs after 561 activation of Chrimson in VA neurons.

##### Trafficking of GFP::UNC-9 particles

L4 larvae were immobilized using 10% agarose pads, 50mM muscimol and 0.5 µm Polybead® Carboxylate Microspheres (Polysciences). VA::GFP::UNC-9 was captured using 488 nm excitation 5 frames/sec for three-minutes on a Nikon Spinning Disk microscope with a 100X SR Apo TIRF (1.49 NA) objective. Movies were obtained from a single plane and were deconvolved with Nikon 2D deconvolution software. A 5-pixel line was drawn through the VA process to create a kymograph in Nikon elements. The kymograph was then analyzed using KymoButler Premium. The number of GFP::UNC-9 puncta was determined based on fluorescence intensity and size. Puncta were considered motile if they moved at a speed > 0.4 µm/sec for three consecutive frames. The percentage of puncta that moved in a given VA were calculated. Puncta velocity, direction, and displacement were calculated for each punctum with directed locomotion. A Mann-Whitney was used to determine significance between groups.

##### Fluorescence Recovery After Photobleaching (FRAP) of GFP::UNC-9 particles

Individual L3 larvae were placed on 10% agarose pads and immobilized with 50mM muscimol. A single anterior VA neuron (VA2, VA3, or VA4) in each experimental animal was imaged with 488 nm excitation to collect a Z-stack (0.2 µm/step) on a Nikon Spinning Disk with a 100X SR Apo TIRF (1.49 NA) objective. A 405 nm laser (15% power, 15ms dwell time) was used to bleach an ROI encompassing the anteriorly directed VA axon but excluding the VA cell soma which was not bleached. An additional Z-stack with 488 nm excitation was captured immediately after photobleaching. The treated worm was recovered by washing the slide with M9 and allowed to recover on a bacterially seeded (OP50-1) NGM plate until reaching the adult stage (24 hours at 20°C). Each animal was then placed on a 10% agarose pads and immobilized with 50mM muscimol. GFP::UNC-9 signal was imaged from the previously photobleached region of each treated VA neuron. GFP::UNC-9 signal was summed from a 5-pixel-wide line drawn along the VA process. The total amount of corrected fluorescence (background subtracted) was measured and divided by the length of the process to account for growth of the VA process during development. The resultant GFP::UNC-9 density (Total fluorescence/10 µm) was normalized to the GFP::UNC-9 density at the initial timepoint at the L3 larval stage before photobleaching. A paired t-test was used to determine significance between groups.

### QUANTIFICATION AND STATISTICAL ANALYSIS

For all categorical data, we used a Fisher’s exact test. For all quantitative data, we used Prism9 to determine if a sample was normally distributed. For normally distributed samples, a Student’s t-test (2 groups) or one-way ANOVA with multiple comparisons correction (3 or more groups) was used. If a sample in a given analysis was not normally distributed, a Mann-Whitney (2 groups) or Kruskal-Wallis test (3 or more groups) was used. Figure legends specify the statistical test and N used in each experiment.

### KEY RESOURCES TABLE

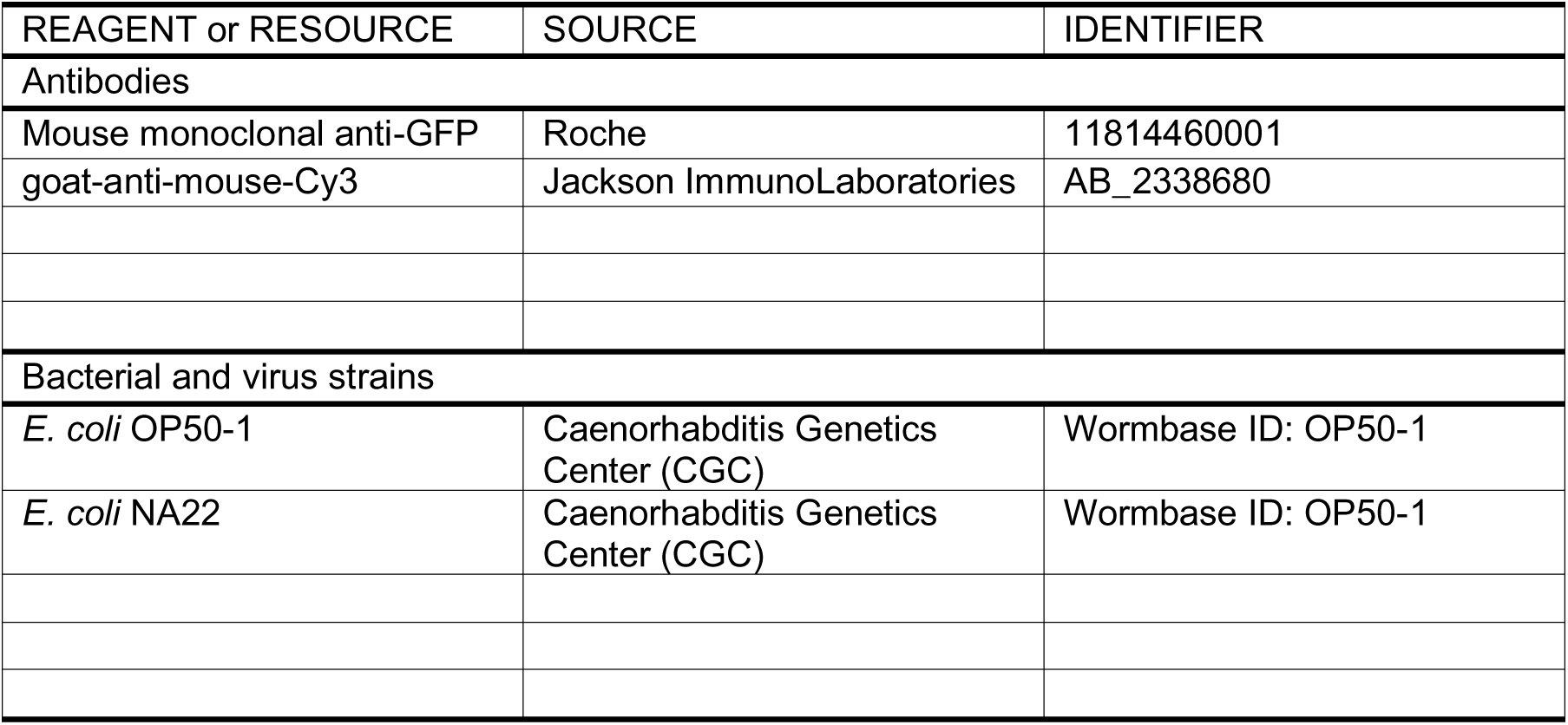
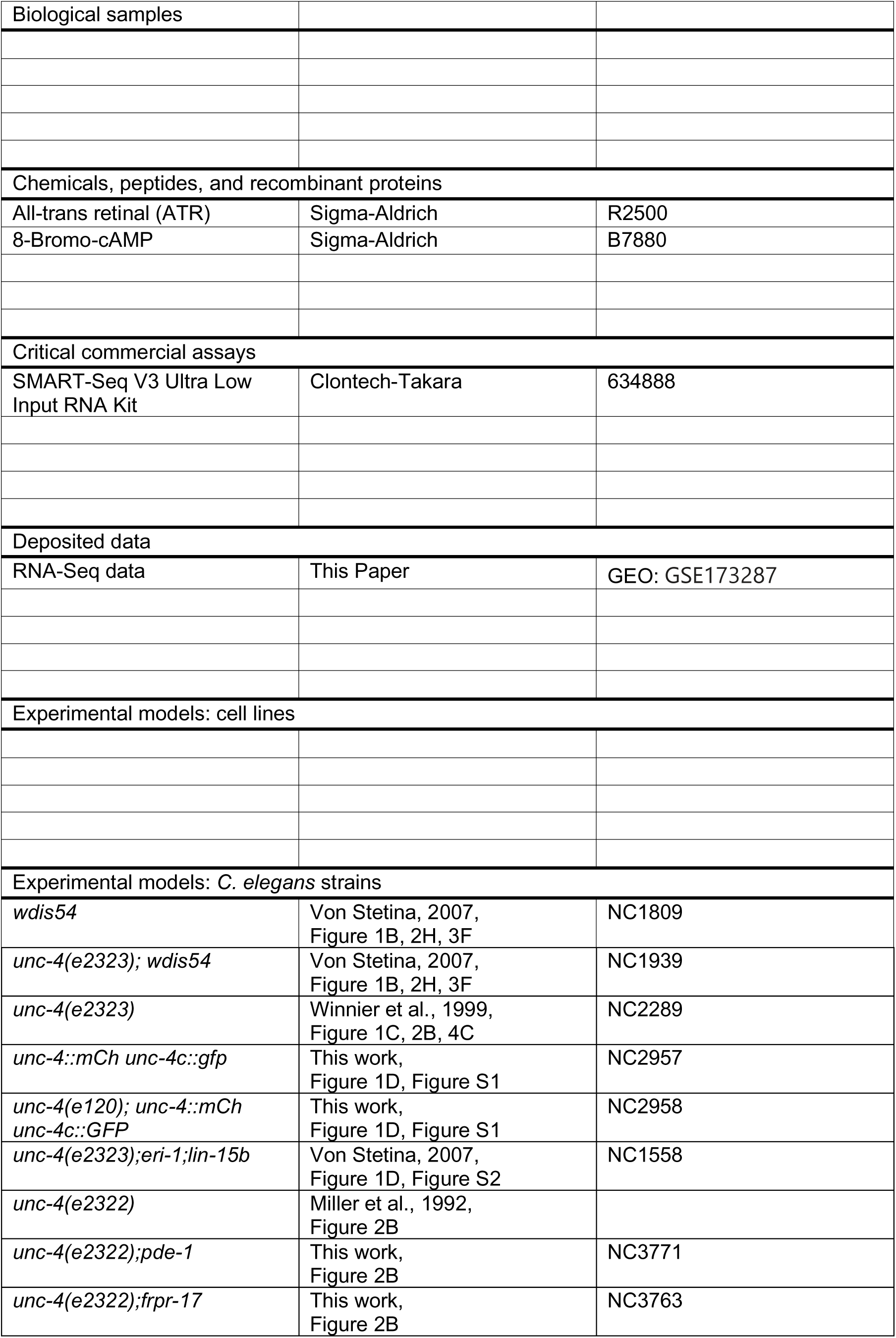
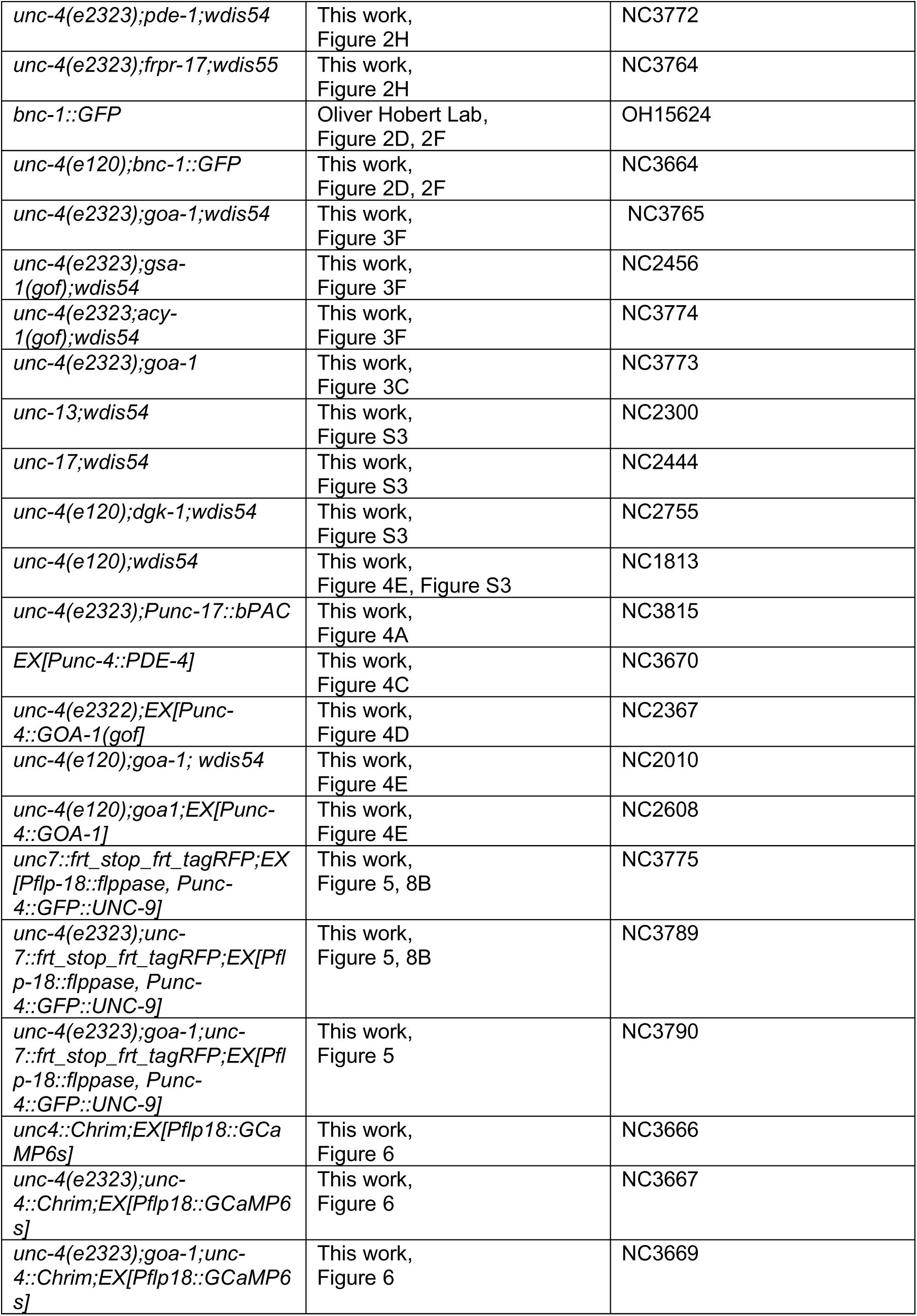
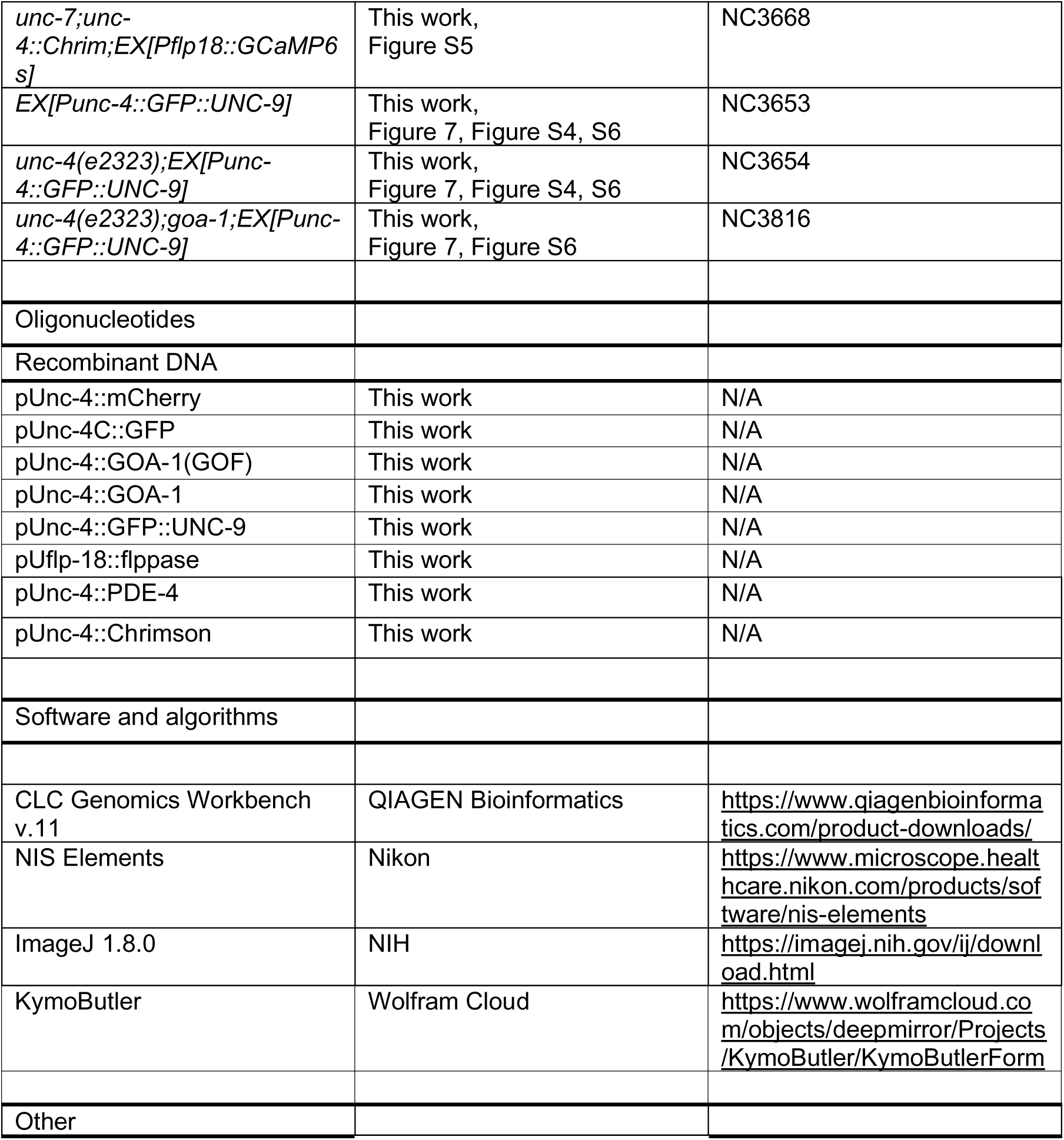

