## Supplemental Files for "cAMP controls a trafficking mechanism that directs the neuron specificity and subcellular placement of electrical synapses"

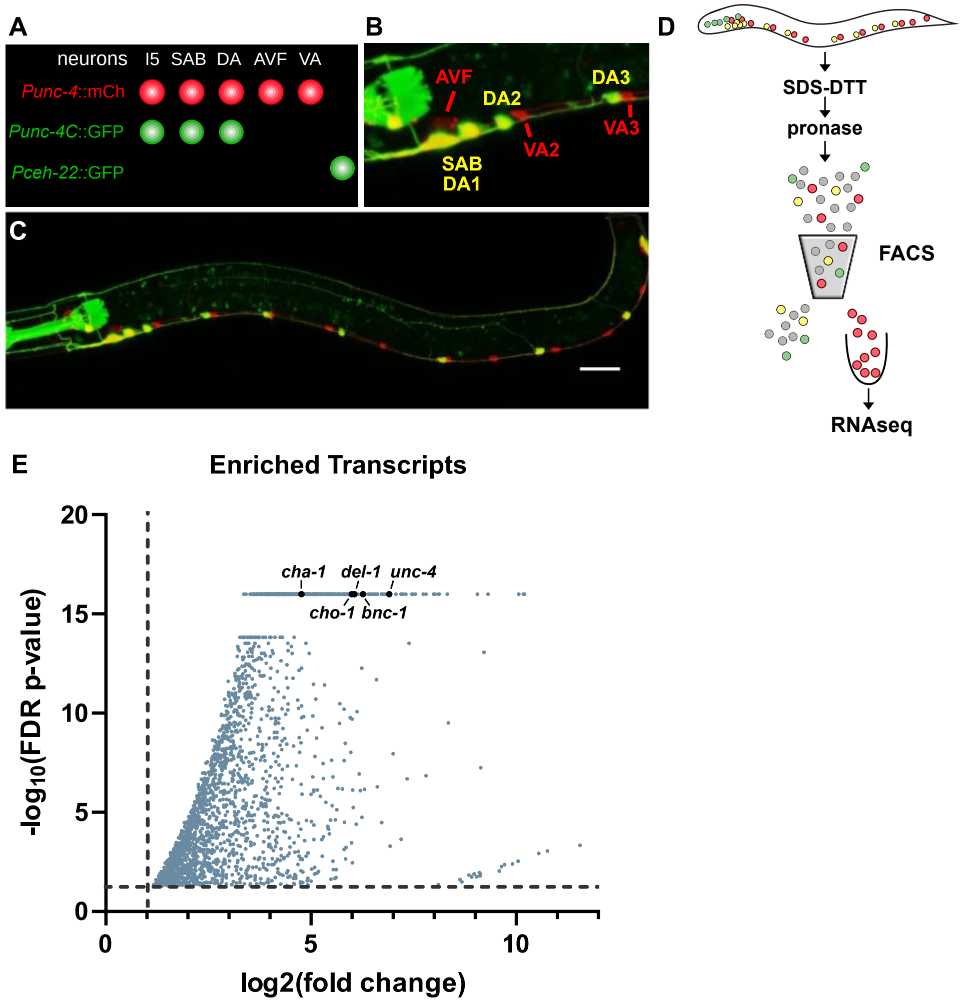

### Supplemental Figure 1: Strategy to detect VA enriched transcripts

**A.** Diagram representing neurons labeled with intersectional strategy to enrich for VA motor neurons. The full length *unc-4* promoter driving mCherry labeled I5, SABs, DAs, AVFs, and VA motor neurons in red. The truncated *unc-4C* promoter was used to label I5, SABs, and DAs with green. As a result, only VAs and AVFs were solely labeled in red. As VA motor neurons outnumber AVFs 12:2, we reasoned that this strategy would predominantly label VA motor neurons.
**B.** Zoomed in image of anterior portion of a worm labeled with intersectional strategy. VAs are labeled in red, as are AVFs (dim red).
**C.** Image of full worm expression intersectional fluorophores to enrich for VA motor neurons.
**D.** Schematic representation of pipeline to isolate VA motor neurons for RNA-seq. Labeled worms were synchronized to capture a large number of L2 worms which were then dissociated to single cells and subjected to FACS for collection of red-only (VA enriched) cells. RNA was isolated from the FACS cells and subjected to RNA-seq.
**E.** Volcano plot of upregulated transcripts (> 2X, FDR p-value < .01) detected in wild type VAs (see Methods). Characteristic genes of VA motor neurons were of the most significantly upregulated genes detected (*cha-1, del-1, unc-4, cho-1, bnc-1*).

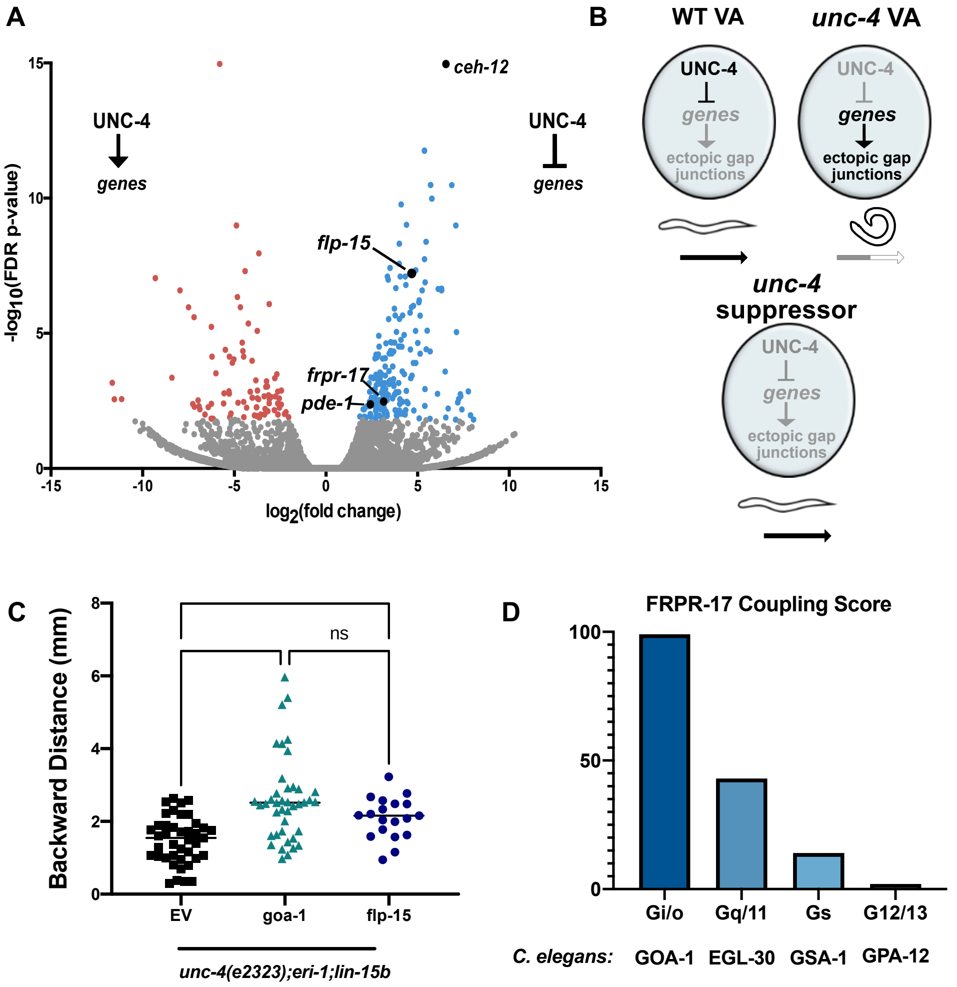

**Supplemental Figure 2: Identifying suppressors of the Unc-4 movement defect**

**A.** Volcano plot of upregulated transcripts (> 2X, FDR p-value < .01) (blue dots) and downregulated (< -2X, FDR p-value <.01) (red dots) detected in *unc-4* mutant VAs compared to wild type VAs (see Methods). Black dots indicate identified suppressors of the Unc-4 movement defect.

**B.** Schematic representation of strategy to identify potential regulators of gap junction specificity in Unc-4 "suppressor" screen. In wild type VAs (**top left**) UNC-4 is expressed and prevents the expression of genes that lead to ectopic gap junctions and loss of backward movement. In *unc-4* mutant VAs (**top right**), UNC-4 is lost and as a result, genes that promote ectopic gap junction formation are expressed and these worms cannot move backward. In an *unc-4* "suppressor" VA (**bottom**), individual genes are knocked out or knocked down in an *unc-4* sensitized background. If the gene knocked down is required for formation of ectopic gap junctions, miswiring should be partially lost and backward locomotion should be partially restored.

**C.** Loss of flp-15, partially restores backward locomotion to *unc-4* mutant. Quantification of backward distance traveled by *unc-4(e2323);eri-1;lin-15b* fed RNAi of empty vector (black) (n = 41), *goa-1* (teal) (n=41), or flp-15 (blue) (n=20) in a 3-minute period. One-way ANOVA, * p = .0368, **** = p < .0001. Data are mean +/- SE.

**D.** Results of PREDCouple2 prediction of FRPR-17 G-protein coupling. FRPR-17 was predicted (Score 99) to couple to G-proteins that are Gi/o, with lower scores for other classes of G-proteins. The correlating C. elegans G-proteins are listed below.

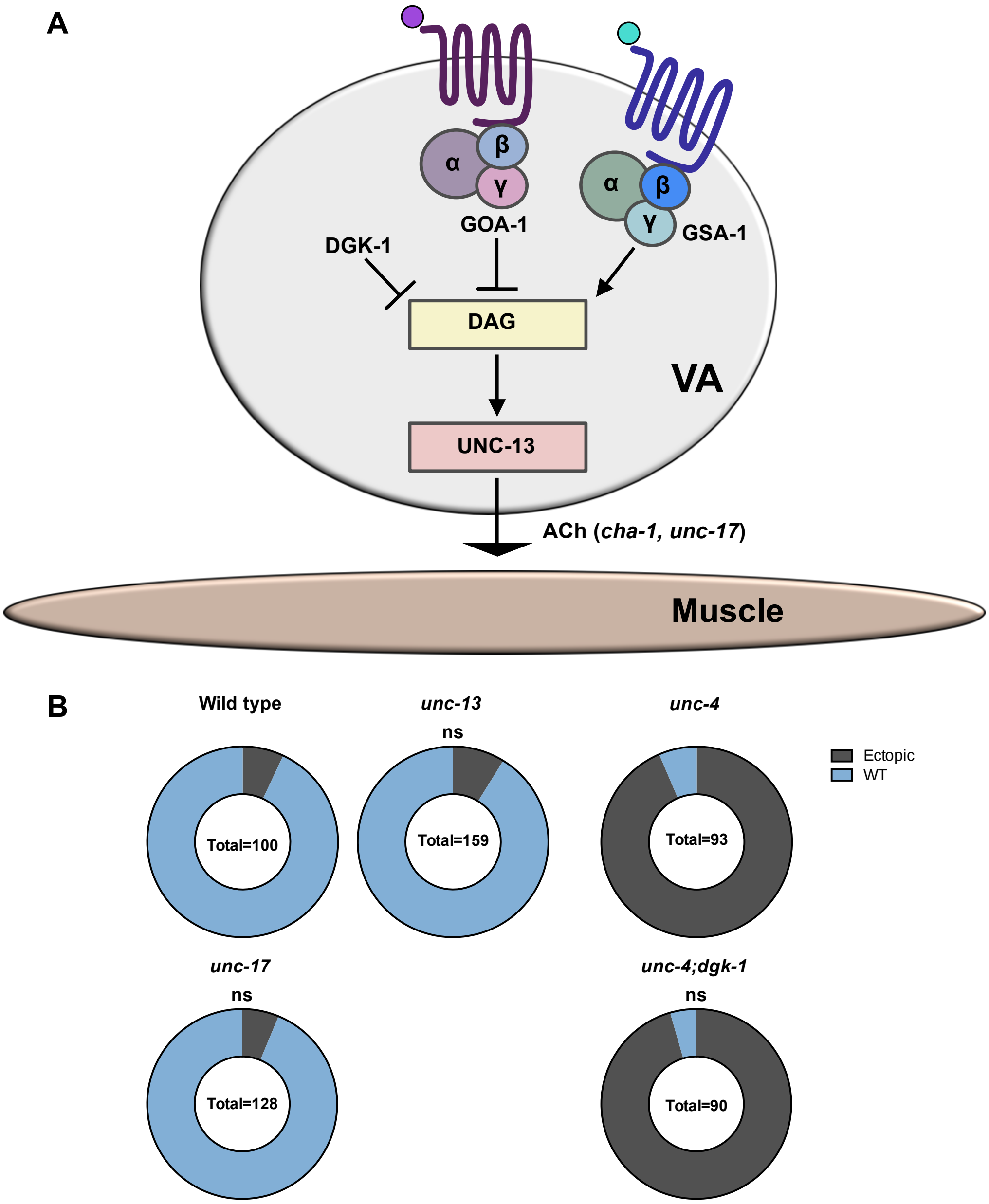

**Supplemental Figure 3: Cholinergic release at the Neuromuscular Junction does not affect gap junction specificity of VAs**

**A.** Schematic of acetylcholine (Ach) release onto muscle which is regulated by GOA-1 and GSA-1. GOA-1 antagonizes DGK- 1/diacylglycerol kinase to inhibit DAG production. GSA-1 functions through ACY-1/adenylyl cyclase and cAMP to promote DAG binding to UNC-13, which is required for synaptic vesicle fusion with syntaxin. As a result, GOA-1 signaling inhibits Ach release while GSA-1 promotes Ach release. Loss of *unc-13* or *unc-17* should result in reduced Ach signaling while loss of *dgk-1* should result in upregulated Ach signaling.

**B.** (**Left**) Percentage of VAs miswired in wild type, *unc-13(e51)* mutants and *unc-17(e113)* mutants. Loss of neither *unc-13* nor *unc-17* results in ectopic miswiring of VA→AVB gap junctions. (**Right**) Percentage of VAs with ectopic VA→AVB gap junctions in *unc-4(e120)* mutants, and *unc-4;dgk-1* double mutants.  Loss of *dgk-1* does not suppress the *unc-4* miswiring defect.

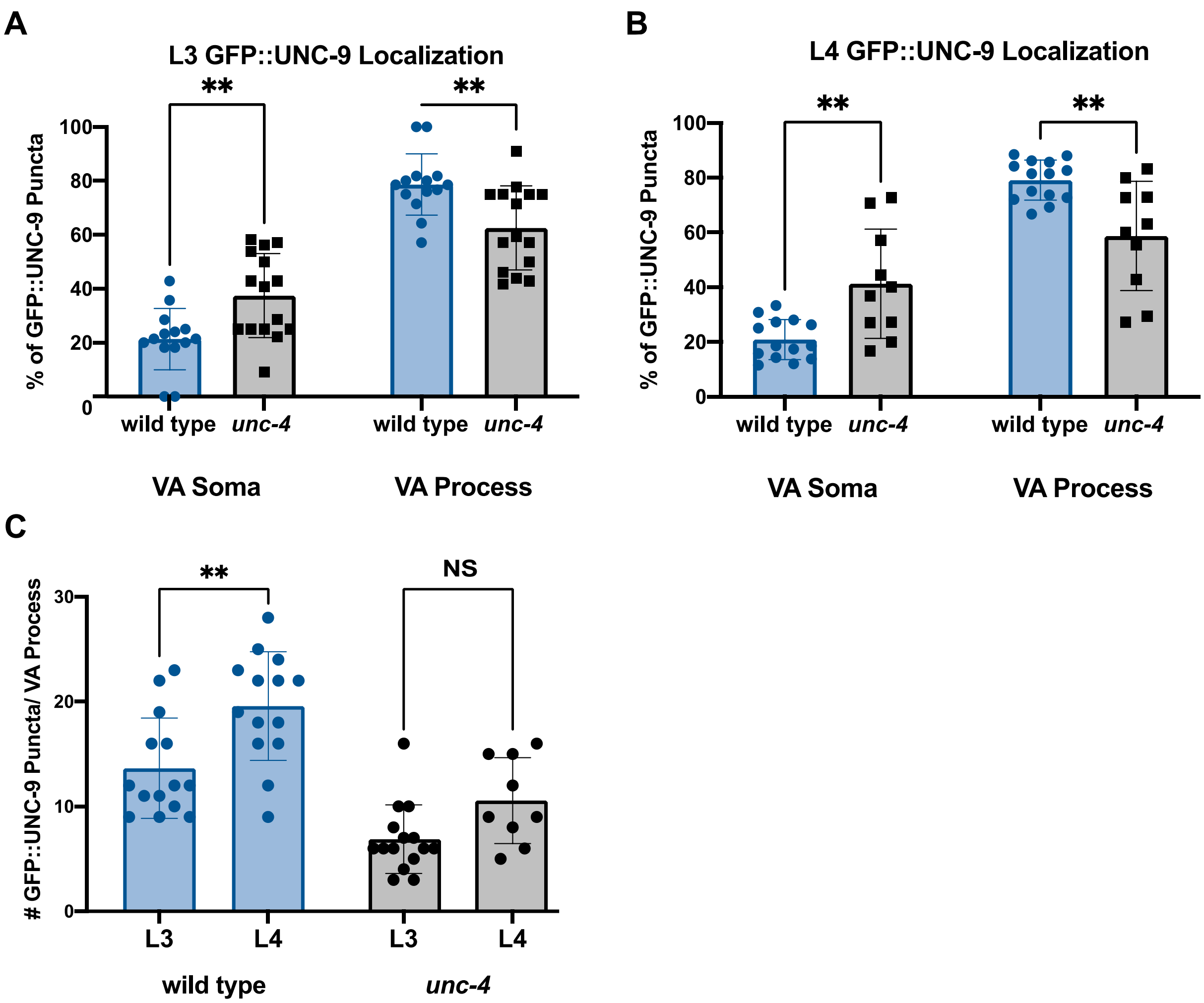

**Supplemental Figure 4: GFP::UNC-9 is localized as predicted in VA motor neurons**
**A.** Subcellular localization of *punc-4::GFP::UNC-9* in VA motor neurons at the L3 stage in wild type (blue) and *unc-4* mutants (black). *Unc-4* mutant VAs have significantly more GFP::UNC-9 puncta on the cell soma and significantly fewer puncta within the VA process compared to wild type, as predicted by EM reconstructions (White et al., 1986, 1992). Two-way ANOVA used to determine significance. ** = $p < .001$. N = 15 for each group.
**B.** Subcellular localization of *punc-4::GFP::UNC-9* in VA motor neurons at the L4 stage in wild type (blue) and *unc-4* mutants (black). *Unc-4* mutant VAs have significantly more GFP::UNC-9 puncta on the cell soma and significantly fewer puncta within the VA process compared to wild type. Two-way ANOVA used to determine significance. ** = $p < .001$. N >10 for each group.
**C.** The number of GFP::UNC-9 puncta in the VA process over time in wild type (blue) and *unc-4* (black). In wild type worms, significantly more GFP::UNC-9 puncta can be detected in the VA process at the L4 stage compared to the L3 stage. This increase in GFP::UNC-9 number is not observed in *unc-4.* Two-way ANOVA used to determine significance. ** = $p < .001$. NS = Not Significant. N >10 for each group.

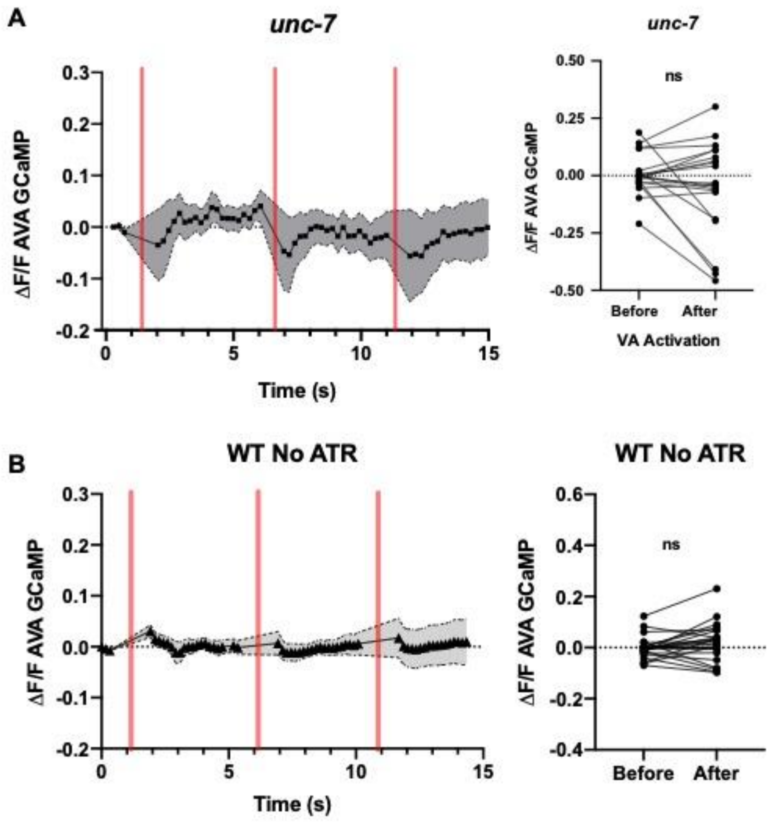

**Supplemental Figure 5: Detection of VA→AVA electrical synapses**
**A.** VA→AVA electrical synapses are not detected in *unc-7* mutants in which VA→AVA gap junctions are lost (Starich et al., 2009). (**left**) Quantification of *unc-7* AVA::GCaMP6s $\Delta F/F_0$ of *AVA::GCaMP6s* fluorescence over time. Three successive VA activations (500 ms) are denoted by red vertical bars. Shaded area = SEM. N = 9 worms. (**Right**) Quantification of $\Delta F/F_0$ before versus after 561 stimulation. Paired t-test. N = 9 worms, 18 activations. NS = Not Significant.
**B.** AVA GCaMP response is dependent on VA activation. Calcium influx in AVA upon VA activation is not detected in wild type womrs grown in the absence of ATR, the necessary cofactor of Chrimson. (**left**) Quantification of wild type (-ATR) AVA::GCaMP6s $\Delta F/F_0$ of *AVA::GCaMP6s* fluorescence over time. Three successive VA activations (500 ms) are denoted by red vertical bars. Shaded area = SEM. N = 9 worms. (**Right**) Quantification of $\Delta F/F_0$ before versus after 561 stimulation. Paired t-test. N = 9 worms, 18 activations. NS = Not Significant.

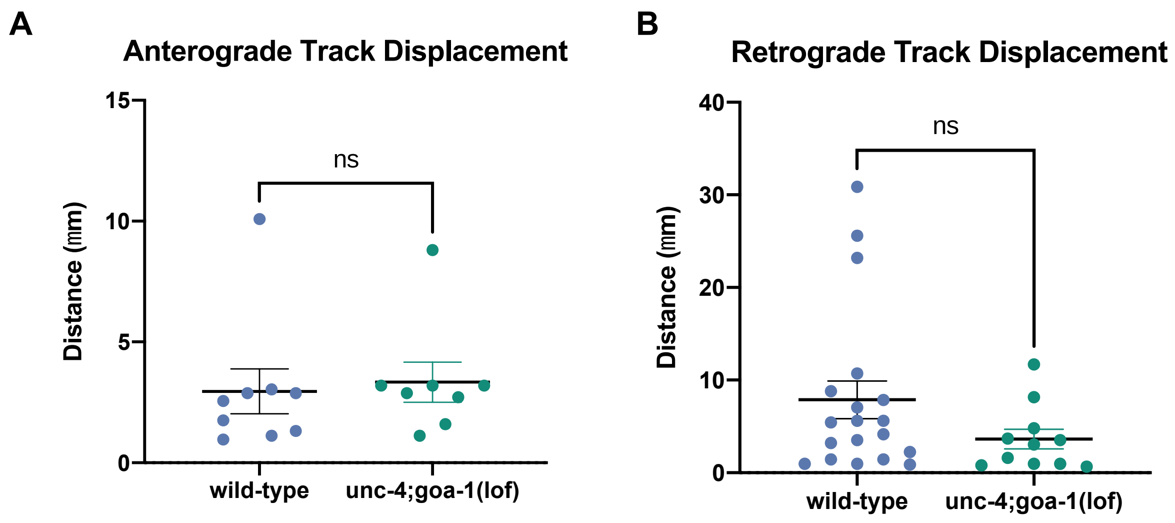

**Supplemental Figure 6: Track displacement of GFP::UNC-9**

**A)** Quantification of the anterograde track displacement of individual GFP::UNC-9 puncta in wild-type (Mean = 2.956 um, SEM=.9039) and *unc-4;goa-1(lof)* (Mean = 3.340 um, SEM=.8283 VA motor neurons. N=9 for each genotype. Student's t-test, NS= not significant.

**B)** Quantification of the retrograde track displacement of individual GFP::UNC-9 puncta in wild-type (Mean = 7.869 um, SEM=2.036) and *unc-4;goa-1(lof)* (Mean = 3.622um, SEM=1.055) VA neurons. N=20 for wild type and N=10 for *unc-4;goa-1*. Student's t-test, NS= not significant.

**Supplemental Table 1:** Genes tested for suppression of Unc-4 backward locomotion

| **Gene Tested for Backward Suppression** | **Method** | **Suppressed?** |
|---|---|---|
| *acr-15* | RNAi | |
| *asah-1* | RNAi | |
| *C30B5.7* | RNAi | |
| *C30F12.5* | RNAi | |
| *C34C6.7* | RNAi | |
| *C35E7.2* | RNAi | |
| *C35E7.4* | RNAi | |
| *C47D2.1* | RNAi | |
| *ccb-2* | RNAi | |
| *ceh-12* | Mutant and RNAi | Yes |
| *ceh-24* | RNAi | |
| *ceh-31* | RNAi | |
| *ces-1* | RNAi | |
| *ckr-1* | Mutant | |
| *cof-2* | RNAi | |
| *cup-4* | RNAi | |
| *D2024.4* | RNAi | |
| *exc-5* | RNAi | |
| *F10B5.3* | RNAi | |
| *F11E6.6* | RNAi | |
| *F14F11.2* | RNAi | |
| *F26B1.1* | RNAi | |
| *F32f2.1* | RNAi | |
| *F34D10.4* | RNAi | |
| *F42A8.1* | RNAi | |
| *F43G6.4* | RNAi | |
| *F44G4.7* | RNAi | |
| *flp-1* | Mutant | |
| *flp-10* | RNAi | |
| *flp-15* | RNAi | Yes |
| *flp-9* | RNAi | |
| *frpr-15* | RNAi | |
| *frpr-17* | Mutant and RNAi | Yes |
| *gcy-11* | RNAi | |
| *gcy-15* | RNAi | |
| *gcy-21* | RNAi | |

| | | |
|---|---|---|
| *glr-2* | RNAi | |
| *hil-7* | RNAi | |
| *K01A2.3* | RNAi | |
| *K01A6.6* | RNAi | |
| *K07D4.5* | RNAi | |
| *lec-2* | RNAi | |
| *lim-4* | RNAi | |
| *lim-7* | RNAi | |
| *lin-12* | RNAi | |
| *lin-31* | RNAi | |
| *nlp-12* | RNAi | |
| *nlp-15* | RNAi | |
| *nlp-38* | RNAi | |
| *nlp-40* | RNAi | |
| *pax-2* | RNAi | |
| *pct-1* | RNAi | |
| *pde-1* | Mutant | Yes |
| *prkl-1* | Mutant | |
| *R05G6.10* | RNAi | |
| *R06C1.6* | RNAi | |
| *rgs-3* | RNAi | |
| *sem-4* | RNAi | |
| *ser-5* | Mutant | |
| *sod-4* | RNAi | |
| *sue-1* | RNAi | |
| *T19C3.5* | RNAi | |
| *T23G7.3* | RNAi | |
| *unc-122* | RNAi | |
| *unc-129* | RNAi | |
| *unc-30* | RNAi | |
| *unc-71* | RNAi | |
| *unc-86* | RNAi | |
| *vab-23* | RNAi | |
| *vab-8* | Mutant | |
| *W04A8.4* | RNAi | |
| *Y43C5A.3* | RNAi | |
| *Y47H9A.1* | RNAi | |
| *ZC581.3* | RNAi | |
| *zig-5* | RNAi | |

| | |
|---|---|
| *zip-8* | RNAi |
| *ZK265.7* | RNAi |

**Supplemental Table 2: High prevalence of adjacent neurons that fail to form electrical synapses.** Tab 1: List of pairs of neurons that form chemical synapses but not electrical synapses. This list is likely an underestimate of neurons that physically contact but don't form electrical synapses. Tab 2: Neurons from Tab 1 color coated by expression of the innexins UNC-7 and UNC-9. Over 600 neurons express compatible innexins, physically contact, and do not form gap junction.

**Supplemental Video 1: *unc-4* mutants exhibit backward locomotion defect by tapping assay**

**Supplemental Video 2: Super resolution microscopy reveals coincident UNC-7::tagRFP and GFP::UNC-9 are closely localized, but slightly displaced**

**Supplemental Video 3: Evoked AVA GCaMP response. Individual AVA process expressing GCaMP of wild type, *unc-4,* and *unc-4;goa-1* double mutant. VAs expressing Chrimson are evoked and the two subsequent frames are noted with a red circle in the left-hand corner. In wild type and *unc-4;goa-1* worms, a detectable AVA GCaMP response is detected following VA activation**

**Supplemental Video 4: GFP::UNC-9 trafficking in VA process in wild type and *unc-4* mutant.**
